# The majority of axonal mitochondria in mammalian neurons lack mitochondrial DNA and do not produce ATP

**DOI:** 10.1101/2024.02.12.579972

**Authors:** Yusuke Hirabayashi, Tommy L. Lewis, Yudan Du, Emiliano Zamponi, Parker Kneis, Jennifer U. Jones, Aubrianna M. Decker, Giovanna Coceano, Jonatan Alvelid, Momone Kikuchi, Masafumi Tsuboi, Shogo Suga, Koyo Shibayama, Maëla A. Paul, Daniel M. Virga, Stevie Hamilton, Abigail Morgan, Michael J Beckstead, Pina Colarusso, Timothy E. Shutt, Yasufumi Takahashi, Jellert T. Gaublomme, Ilaria Testa, Franck Polleux

**Author notes:** Co-corresponding authors: Franck Polleux, Columbia University, Department of Neuroscience, Mortimer B. Zuckerman Mind Brain Behavior Institute, Jerome L. Greene Science Center, 3227 Broadway, L-5-050, MC 9853, New York, NY 10027, USA, Yusuke Hirabayashi, Tommy L Lewis Jr. These authors contributed equally to this work (Yusuke Hirabayashi, Tommy L. Lewis, Yudan Du, Emiliano Zamponi).

## Abstract

In neurons of the mammalian central nervous system (CNS), axonal mitochondria are thought to be indispensable for supplying ATP during energy-consuming processes such as neurotransmitter release. We have previously shown that glutamatergic excitatory cortical pyramidal neurons (CPNs) are characterized by a striking compartmentalization of mitochondrial structure. Axonal mitochondria are maintained at a small uniform size (∼1 microns length) by high levels of MFF-dependent fission, whereas dendritic mitochondria form a highly fused network of mitochondria throughout the entire dendritic arbor(1, 2). Here, we tested whether these striking structural differences reflect functional and molecular compartmentalization. Using four independent techniques, we reveal that in CPNs as well as in two subtypes of GABAergic inhibitory cortical interneurons and neuromodulatory dopaminergic neurons of the substantia nigra, the majority of axonal, but not dendritic, mitochondria lack mitochondrial DNA (mtDNA) *in vivo*. We also show that axonal mitochondria are depleted in both mtDNA-encoded mRNAs and proteins compared to dendritic mitochondria. Timelapse imaging establishes that most mitochondria entering nascent dendritic and axonal processes from the soma are small and lack mtDNA. Using dynamic, optical imaging of genetically encoded sensors for ATP and pH targeted to the mitochondrial matrix, we demonstrate that in axons of CPNs, but not in their dendrites, mitochondrial F_1_F_0_-ATP synthase (Complex V) functions in a reverse way, hydrolyzing ATP and extruding H^+^ out of the matrix to maintain mitochondrial membrane potential. Our results indicate that in neurons of the mammalian CNS, the majority of axonal mitochondria lack mtDNA and likely do not play a major role in ATP generation, despite playing other functions such as regulation of neurotransmission via presynaptic Ca^2+^ buffering(1, 3–5).

---

Mitochondria are often referred to as the ‘powerhouse’ of the cell because of their ability, through oxidative phosphorylation, to generate large amounts of ATP. However, mitochondria play many other critical functions such as regulating cytosolic Ca^2+^ dynamics and lipid biogenesis (6). The human brain is often considered the most energy consuming organ in our body representing only 2% of our body mass but consuming up to ∼20% of glucose (7). In neurons of the central nervous system (CNS), mitochondria-dependent ATP synthesis through oxidative phosphorylation is thought to play a significant role in supporting key, energy-consuming, neuronal functions such as presynaptic neurotransmitter release along axons (8–13).

However, several observations cast doubt on how critical axonal mitochondria are for ATP generation through oxidative phosphorylation (OxPhos) in mammalian CNS neurons: (1) only 50% of presynaptic boutons along axons of CPNs are associated with mitochondria and these mitochondria are uniformly small in size (∼1μm length) (1, 3, 14, 15), (2) presynaptic release sites lacking mitochondria are characterized by higher neurotransmitter release probability than presynaptic boutons associated with mitochondria (3, 4), and (3) blocking oxidative phosphorylation in mammalian neurons has limited effects on presynaptic ATP concentration, even under extreme, non-physiological, levels of action potential-triggered stimulation of presynaptic release (11, 16). Of note, a recent proteomic study using synaptosomes isolated from glutamatergic CPNs revealed an enrichment in glycolytic proteins and a relative depletion in proteins involved in oxidative phosphorylation, a trend opposite to what is observed in synaptosomes isolated from GABAergic interneurons such as parvalbumin (PV)-positive fast-spiking interneurons (17). Surprisingly, cell type-specific, conditional, deletion of mtDNA-associated protein Twinkle has significant phenotypic consequences on astrocyte maintenance but relatively minor effects on neuronal survival until 8 months after birth, despite loss of mtDNA in neurons, arguing that neurons can tolerate mtDNA loss far better than astrocytes in the mammalian CNS (18). Lastly, alternate sources of ATP generation are available to neurons *in vivo*, such as glycolysis which is highly functional in axons (7, 19–23).

Since the mitochondrial genome contains 13 protein-coding genes essential for all complexes of the electron transport chain (ETC) required for OXPHOS, we first assessed the presence of mtDNA in individual dendritic versus axonal mitochondria in developing and mature mouse CPNs *in vitro* and *in vivo*. Using multiple, independent *in vitro* and *in vivo* approaches including (1) immunofluorescence detection of mtDNA and visualization of mtDNA-associated proteins such as Twinkle and TFAM, (2) single molecule DNA-FISH for mtDNA, as well as (3) quantitative PCR detection of mtDNA from single axonal mitochondria isolated using Scanning Ion Conductance Microscopy (SICM), our results demonstrate that mtDNA is undetectable in ∼80-90% of axonal mitochondria of mouse cortical CPNs, while detected in 60-80% of dendritic mitochondria of the same neurons. Quantification of mtDNA content in axonal mitochondria of three other neuronal subtypes, both somatostatin (SST)- and PV-positive cortical GABAergic inhibitory interneurons, as well as dopaminergic (DA) neurons from the substantia nigra pars compacta (SNc) reveals similar ranges (70-90%) of mtDNA-negative axonal mitochondria. Finally, live imaging analysis using mitochondrial matrix-targeted, genetically encoded sensors for ATP (mt-iATPSnFR1.0) as well as pH (mt-SypHer) demonstrate that in axonal, but not in dendritic mitochondria, complex V (ATP synthase) functions in a reverse way, hydrolyzing ATP and extruding H^+^ out of the matrix to maintain mitochondrial membrane potential. Together, our results suggest a major revision of the role of axonal mitochondria, at least in mammalian neurons of the CNS. We find that axonal mitochondria are unlikely to play a major role in ATP generation, despite playing other critical and previously characterized functions at presynaptic release sites such as Ca^2+^ buffering to regulate neurotransmitter release probability (1, 3–5).

## Most axonal mitochondria lack mtDNA and mtDNA-associated proteins in CPNs *in vitro* and *in vivo*

We used *ex utero* electroporation (EUE), performed at E15 to target progenitors generating layer 2/3 CPNs, to sparsely express the fluorescently tagged mtDNA-associated protein Twinkle (Twinkle-Venus) and an outer mitochondrial membrane (OMM) targeted mCherry (mCherry-ActA) (**Fig. 1**), followed by dissociation and maintenance in high-density cultures for 5-15 days *in vitro* (DIV). These high-density neuronal cultures contain both cortical neurons (only cell type expressing the plasmids mentioned above) as well as non-neuronal cells such as astrocytes that are critical for neuronal survival and synapse formation. Upon fixation, we coupled fluorescent detection of Twinkle-Venus and mCherry-ActA with an antibody-based detection of DNA (24) (**Fig. 1A**). The anti-DNA signal present outside the neurons co-expressing Twinkle-Venus and mCherry-labelled mitochondria originate from the unlabeled neurons and astrocytes present in the culture.

**Figure 1.**
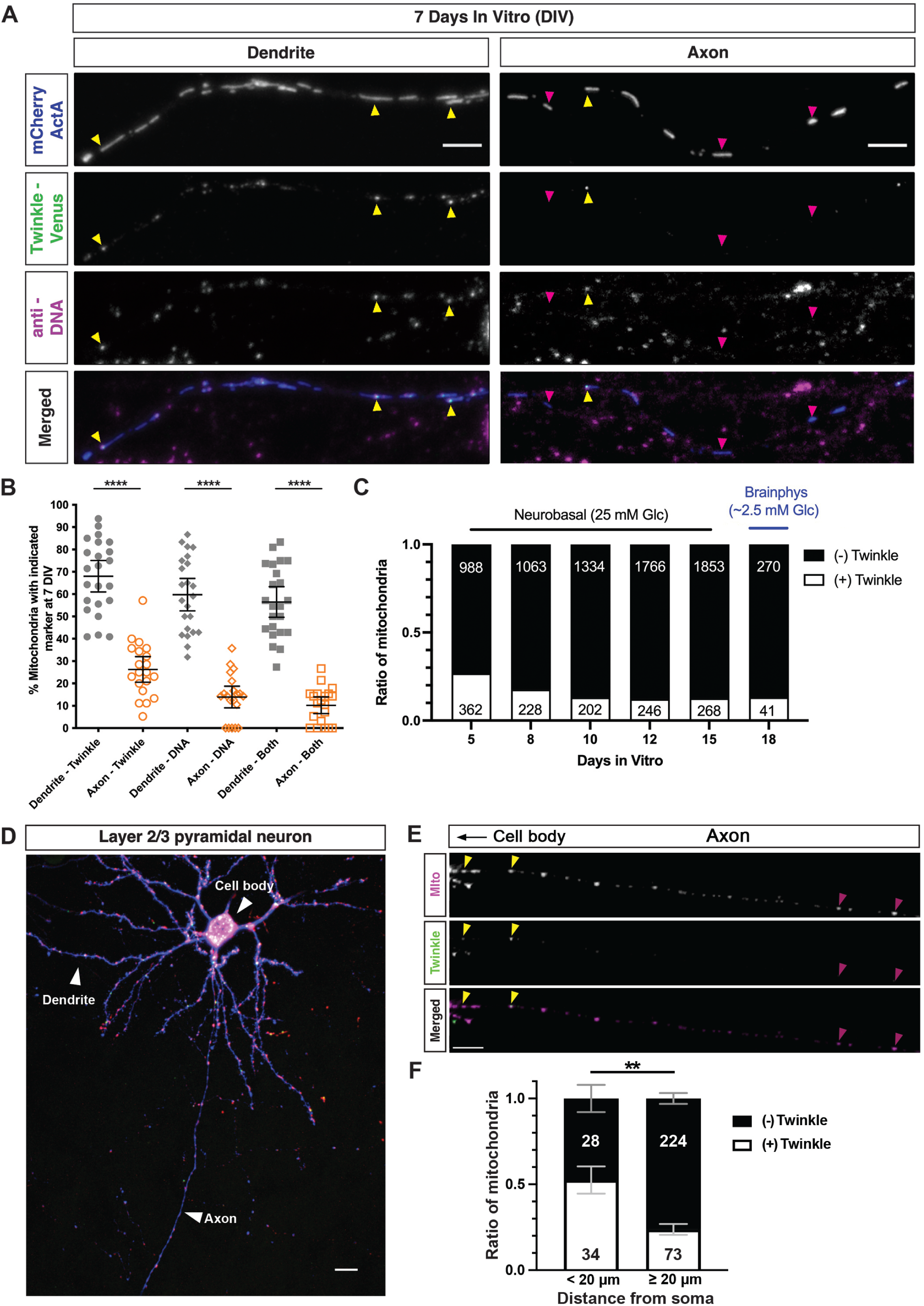
A low fraction of axonal mitochondria contain mtDNA and mtDNA-associated proteins compared to dendritic mitochondria of cortical pyramidal neurons *in vitro* and *in vivo*. (**A**) Representative dendritic (left) or axonal (right) segments from CPNs at 7DIV expressing mCherry-ActA (blue) to visualize mitochondria, Twinkle-Venus (green) to visualize the mitochondrial nucleoid and stained with an anti-DNA antibody (magenta) to directly visualize mtDNA. DNA+/Twinkle+ mitochondria are indicated by yellow arrows and DNA-/Twinkle- mitochondria are indicated by magenta arrows. (**B**) Quantification of the percentage of mitochondria with the indicated marker (or mitochondria positive for both DNA and Twinkle – labeled as Both) in dendrites or axons (each point represents an individual segment). N_dendrite_ = 318 mitochondria from 23 dendrite segments, N_axon_ = 298 mitochondria from 20 axon segments from 3 independent experiments. **** p ≤ 0.0001 by Kruskal-Wallis test. Error bars are 95% confidence intervals of the mean. (**C**) Quantification of the fraction of Twinkle+ axonal mitochondria at various times in culture using Neurobasal containing 25mM glucose (5-15DIV) or in Brainphys containing 2.5mM glucose (see Extended Data Fig. 1H for detail) with mean ± SEM (5DIV neurons: n = 4 independent dishes and 8 or 10 or 12 or 18DIV neurons: n = 3 independent dishes). Numbers of mitochondria are indicated in each column. (**D-E**) Representative images of a layer 2/3 CPNs *in vivo* (D; its axon, dendrites and cell body highlighted by arrowheads) expressing mTagBFP2 (blue), Twinkle - Venus (green) and OMM-targeted mCherry-ActA (red) at P27. The proximal portion of its axon emerging from the cell body is shown at higher magnification (E) with mCherry-ActA pseudocolored in magenta and Twinkle-Venus in green. Yellow arrowheads indicate Twinkle+ mitochondria in proximal part of axon, magenta arrowheads indicate Twinkle- mitochondria in more distal portion of axon. Scale bar: 10 μm. (**F**) Ratio of Twinkle positive (+) or negative (-) mitochondria in the proximal (< 20 μm) or distal (> 20 μm) portion of the axon emerging from the soma. Numbers of mitochondria counted are indicated in each column. 16 neurons from 5 mice were used for quantifications. ** p<0.0001 by Fisher’s exact test.

Quantification of the fraction of mitochondria positive for either one or both DNA immunofluorescence and Twinkle-Venus gives consistent results and reveals striking differences between axons and dendrites (**Fig. 1B**): in dendrites the fraction of mitochondria containing mtDNA is ∼70% but in axons this fraction goes down to ∼10-20%. This low fraction of mtDNA+ and Twinkle+ axonal mitochondria is slightly but significantly higher in immature CPNs *in vitro* (∼25% at 5DIV) and decreases progressively with neuronal maturation down to ∼13% at 10-15DIV (**Fig. 1C**). These neuronal cultures were performed using media containing 25 mM glucose which is commonly used throughout the literature but likely significantly higher than *in vivo* (25, 26). To test if the extracellular concentration of glucose influences the fraction of mitochondria with mtDNA/Twinkle, we performed an additional experiment in low 2.5 mM glucose concentration by progressive dilution, leading to culture conditions where CPNs are maintained for approximately a week in 2.5 mM glucose from 10-18DIV (**Fig. S1H**). Our results demonstrate that the fraction of Twinkle+ axonal mitochondria is still low (13%) in conditions of low extracellular glucose (2.5 mM) and therefore indistinguishable from cultures maintained in 25 mM glucose (**Fig. 1C**).

Using *in utero* electroporation (IUE) at E15 to label neural progenitors generating layer 2/3 CPNs progenitors and examine mature neurons in 1 month old mice *in vivo*, we confirmed that only a low fraction (∼20%) of mitochondria are Twinkle+ along the distal portion of the axon (**Fig. 1D**). Interestingly, we observed that in the portion of axon most proximal to the soma (<20 μm), corresponding to the axon initial segment (AIS; **Fig. S1A-G**), the fraction of Twinkle+ mitochondria is approximately 50%, while the portion of the axon more distal to the soma (>20μm i.e. beyond the AIS) contained a significantly lower fraction of Twinkle+ mitochondria (24%; **Fig. 1D-F**). We obtained the same results in layer 2/3 CPNs *in vivo* using a different nucleoid/mtDNA-associated protein (TFAM-mCherry) expressed by IUE, with less than 20% of axonal mitochondria being TFAM+, whereas ∼60% of dendritic mitochondria are TFAM+ (**Fig. S2**).

In CPNs, many axonal mitochondria are localized at presynaptic release sites (27), but only 50% of presynaptic boutons are associated with mitochondria. We examined whether mtDNA content differed for mitochondria localized at presynaptic boutons or not. To achieve this, we performed EUE to co-express three different reporters in cortical PNs *in vitro*: (1) a mitochondrial matrix-targeted blue fluorescent protein (mito-mTagBFP2), (2) Twinkle-mStayGold (green) to label mtDNA nucleoids, and (3) a presynaptic marker vGlut1-mCherry (red) (**Fig. S2E**). Our results confirmed that ∼55% of mitochondria are associated with presynaptic boutons (**Fig. S2F**) but also demonstrate that a low fraction of mitochondria associated with presynaptic boutons (vGlut1+) or mitochondria not associated with presynaptic boutons (vGlut1-) are Twinkle+ (∼15-20%; **Fig. S2G**).

## The low fraction of axonal mitochondria containing mtDNA is not due to diffraction-limited microscopy

In mammalian cells, mitochondria nucleoids display a uniformly small size on the order of one hundred nanometers (28) and contain both mtDNA and associated proteins such as TFAM and Twinkle (29). As the nucleoid sizes are below the diffraction limit (roughly 200 nm at best for lateral (x,y) resolution in light microscopy), we used a non-diffraction limited approach, customized stimulated emission depletion (STED) nanoscopy, in order to improve our capacity to detect single nucleoids using light microscopy (30). This super-resolution microscopy technique features a 42 ± 9nm (mean+/-SD) lateral resolution (**Figure S3A-C**).

We imaged rat hippocampal neurons maintained for 14-16DIV (**Fig. 2A-E**) and immunolabelled with an antibody detecting the dendritic marker MAP2 (cyan in **Fig. 2C**) and/or the AIS marker, Neurofascin (NF; gray in **Fig. 2E** and **Fig. S3D-E**), as well as an anti-DNA antibody (yellow in **Fig. 2A-C**) and/or anti-TFAM antibody (yellow in **Fig. 2D-E**) to label nucleoids and an antibody against the outer mitochondrial membrane (OMM) protein Tom20 (magenta in **Fig. 2A-C**) or the OMP25 localization peptide (magenta in **Fig. 2D-E**) to label the OMM. Both in MAP2+ dendritic segments (dashed lines in **Fig. 2B**) and in the AIS (dashed lines in **Fig. 2D-E**), the use of STED nanoscopic imaging (**Fig. 2B and 2D**) drastically improved resolution of individual mtDNA+ or TFAM+ nucleoids compared to standard confocal microscopy (**Fig. 2C-E** and **Fig. S3C**).

**Figure 2.**
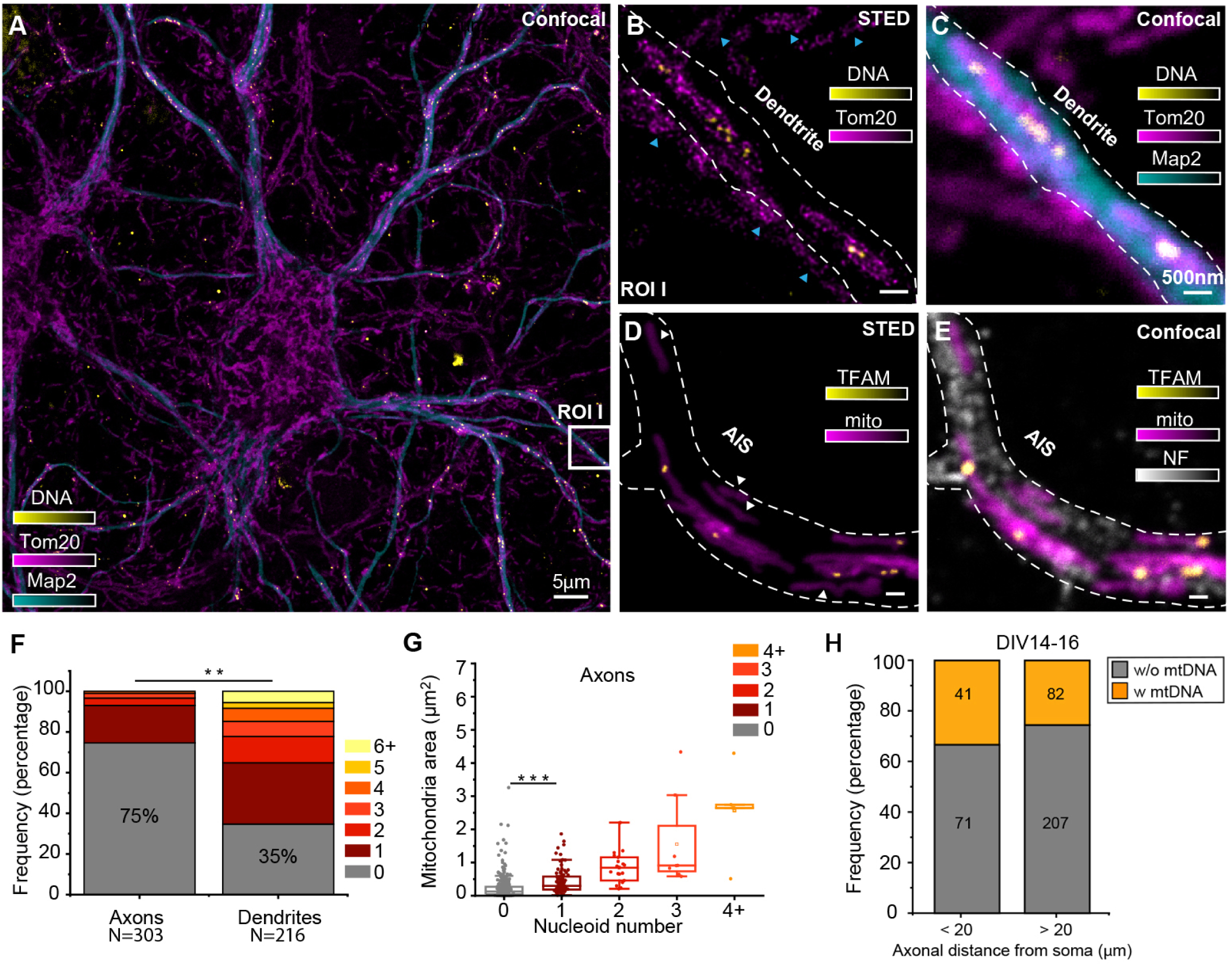
Nanoscopic imaging of mitochondria nucleoids in primary rat hippocampal neurons. **(A)** Large confocal image overview of a neuron in culture showing the organization of mitochondrial DNA in nucleoids; nucleoids are immunostained for DNA (yellow), mitochondria for the outer membrane protein TOM20 (magenta), and dendrites (cyan) for the microtubule associated protein MAP2. **(B)** Two-color STED image of nucleoids from the region-of-interest (ROI) highlighted in A, where single nucleoids are resolved in the STED image, but are unresolved in the confocal comparison (**C**). Nucleoids are not detectable in mitochondria (cyan arrows) outside the dendritic filament (dashed lines), which probably belong to axons wrapping around the dendrite. (**D-E**) Two-color STED image and relative confocal comparison of TFAM (immunostained) and mitochondria (mito) along the axon initial segment (AIS) immunolabelled with an antibody against with the membrane protein Neurofascin (NF). Mitochondria without nucleoids are common in the AIS (white arrows) **(F)** Frequency of mitochondria without (grey) and with (1-6+, colors) nucleoids detected in axons (>20μm from the soma) and dendrites (N_mitochondria_ = 519). ** p < 0.001 by two-sample t-test (t-test). **(G)** Box plot representation of the mitochondria area as a function of nucleoids number in each mitochondrion for axons. **p<0.001 by Kolmogorov-Smirnov test (KS-Test). (**H**) Frequency of mitochondria without (grey) and with (yellow) nucleoids (immunostained for TFAM) detected along the axon (zero at Neurofascin+ AIS emergence from soma) of mature (14-16DIV) rat hippocampal neurons (N_mitochondria_=401).

This improvement in spatial resolution allowed the quantification of the number of nucleoids in axonal and dendritic mitochondria (**Fig. 2F**) as well as the correlation of mitochondria size with nucleoid number per mitochondria in axons (**Fig. 2G**). These results show that the majority (75%) of axonal mitochondria (>20μm away from soma) are devoid of nucleoids detectable by STED microscopy whereas only 35% of dendritic mitochondria lack nucleoids (**Fig. 2F**). Along the axon, we observe a positive correlation between mitochondria size and nucleoid number (**Fig. 2G**). Importantly, quantification of the fraction of nucleoid-containing mitochondria along the length of the axons of rat hippocampal neurons at 14-16DIV (**Fig. 2H**) shows that ∼75% of mitochondria in the distal portion of the axon are TFAM-negative.

## Low fraction of axonal mitochondria with mtDNA in cortical GABAergic interneurons and dopaminergic neurons from the substantia nigra (SNc) *in vivo*

Next, we determined whether the low fraction of mtDNA-containing axonal mitochondria observed in excitatory glutamatergic CPNs *in vitro* and *in vivo* is also detected in other neuronal subtypes in the central nervous system. We first examined DA neurons located in the SNc which form extensive, highly branched axons projections in the striatum (31) (**Fig. 3A**). This neuronal subtype degenerates early and preferentially in Parkinson’s disease and was shown to be vulnerable upon mitochondrial metabolic deficiency (32, 33). We performed stereotactic injection of a Cre-dependent AAV9 expressing mitochondrial matrix-targeted yellow fluorescent protein (mito-YFP) in a dopamine transporter (DAT)-Cre knockin mouse to drive Cre-dependent expression of mito-YFP in dopaminergic neurons of SNc (**Fig. S4A-B**). We then stained mtDNA using an anti-DNA antibody and performed high resolution imaging of mitochondria in axons and dendrites of DA neurons (**Fig. 3B**).

**Figure 3.**
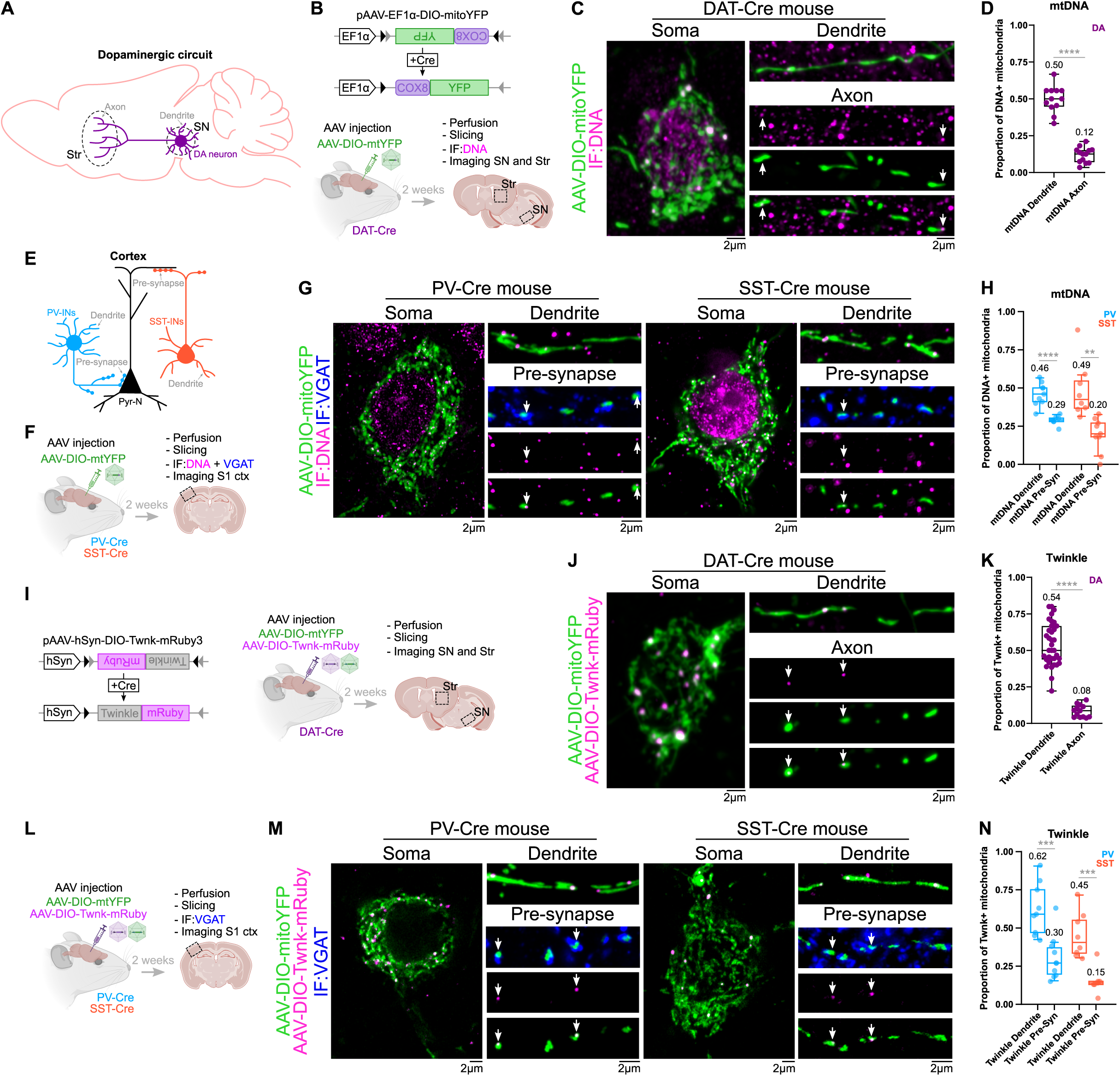
Low fraction of mtDNA-containing mitochondria in axons of dopaminergic neurons, PV+ and SST+ cortical interneurons *in vivo*. (**A**) Schematic representation of a dopaminergic circuit. DA neuron dendritic mitochondria were imaged in the Substantia Nigra (SN) and axonal mitochondria in the Striatum (Str). **(B**) Schematic representation of the construct and experimental design employed to visualize mitochondria and mitoDNA in DA neurons. (**C**) Representative images of mitochondria (green) and mitoDNA puncta (magenta) in the somatic, dendritic, and axonal compartments of DA neurons. Arrows indicate mitochondria containing mitoDNA puncta. (**D**) Quantification of the proportion of DNA-positive mitochondria in dendrites and axons from panel c. Each dot represents individual ROIs, with 169-359 mitochondria from 3-5 mice analyzed per group. p<0.0001 (****) by Mann-Whitney test. (**E**) Schematic representation of a cortical circuit showing PV- and SST-INs making synaptic contacts with the perisomatic and dendritic compartments of cortical pyramidal neurons, respectively. (**F**) Schematic representation of the experimental pipeline utilized to visualize mitochondria and mitoDNA in the dendritic and presynaptic compartment of PV- and SST-INs. (**G**) Representative images of mitochondria (green) with mitoDNA puncta (magenta) in the somatic, dendritic, and presynaptic (VGAT-positive, blue) compartments of PV-and SST-INs. Arrows indicate mitochondria containing mtDNA puncta. (**H**) Quantification of the proportion of DNA-positive mitochondria in the dendritic and presynaptic compartments of PV- and SST-INs from panel g. Each dot represents individual ROIs, with 140-364 mitochondria from 3-5 mice analyzed per group. (**I**) Schematic representation of the construct and experimental approach utilized to visualize mitochondria and Twnk signal in DA neurons. (**J**) Representative images showing mitochondria (green) and Twnk foci (magenta) in the soma, dendrites, and axons from DA neurons. Arrows indicate mitochondria containing mtDNA puncta. (**K**) Quantification of the proportion of Twnk-positive mitochondria in dendrites and axons from DA neurons in panel j. Each dot represents individual ROIs, with 132-359 mitochondria from 3-5 mice analyzed per group. (**L**) Schematic representation of the construct and experimental design utilized to visualize mitochondria and Twnk signal in the dendritic and presynaptic compartment of PV- and SST-INs. (**M**) Representative images showing mitochondria (green) and Twnk puncta (magenta) in the somatic, dendritic, and presynaptic (VGAT-positive, blue) compartments of PV- and SST-INs. Arrows indicate mitochondria containing mtDNA puncta. (**N**) Quantification of the proportion of Twnk-positive mitochondria in the dendritic and presynaptic compartments of PV- and SST-INs from panel m. Each dot represents individual ROIs, with 180-446 mitochondria from 5 mice analyzed per group. p<0.0001 (****), p<0.001 (***) or p<0.01 (**) by t test.

We then examined mtDNA content in axonal and dendritic mitochondria of the two major subtypes of GABAergic (inhibitory) cortical interneurons (INs): (1) PV-INs, also called fast-spiking INs, mediating perisomatic inhibition onto pyramidal neurons (34) and (2) SST-INs, which provide dendrite-targeting inhibition (35) (**Fig. 3E**). Because fast-spiking PV-INs can fire action potentials at very high frequencies (>100Hz) for several seconds, these INs have been reported to have high metabolic demand and specifically high levels of mitochondrial protein expression for components of the electron transport chain (ETC) (36, 37). In fact, a recent proteomic study performed on synaptosomes isolated from genetically labeled neurons including from PV+ interneurons suggested a significant enrichment of proteins involved in oxidative phosphorylation and a relative depletion of glycolytic enzymes, which is the opposite of synaptosomes isolated from glutamatergic CPNs (17).

We labeled mitochondria specifically in PV+ and SST+ cortical INs by stereotactically injecting the same Cre-dependent AAV9 expressing mito-YFP as used above, but in PV-Cre and SST-Cre knockin mice, respectively (**Fig. S4D-F**), followed by DNA immunolabeling and high resolution imaging of axonal and dendritic of PV+ and SST+ cortical INs (**Fig. 3F**). To confidently discriminate axonal mitochondria in cortical GABAergic INs, we specifically focused our analysis on mitochondria localized to the presynaptic compartment, which we labeled using an anti-vesicular GABA transporter (VGAT) antibody. Our results demonstrate that the fraction of mitochondria containing mtDNA in axons of DA SNc neurons and SST+ cortical INs are ∼12% and 20% respectively (i.e. very similar to the values obtained for CPNs *in vitro* and *in vivo* (**Fig. 1** and **Fig. S2**)). Interestingly, we detect a slightly elevated fraction of mtDNA+ axonal/presynaptic mitochondria in PV+ INs reaching ∼30% (**Fig 3H**). In all three neuronal subtypes, we detect significantly higher fraction of dendritic mitochondria containing mtDNA (∼46-50%; **Fig. 3D** and **3H**) compared to their respective axonal mitochondria.

Additionally, we also used an independent approach to label mitochondria and the mtDNA-associated protein Twinkle using two Cre-dependent AAV9 expressing mito-YFP and Twinkle-mRuby fusion protein (**Fig. 3I** and **3L**). Using this approach, we obtained remarkably similar results as with staining for mtDNA (**Fig. 3J-K** and **Fig. 3M-N**): ∼8-15% Twinkle+ in axonal mitochondria in axons of DA SNc neurons and SST+ cortical INs versus 30% Twinkle+ in axonal mitochondria of PV+ cortical INs. Comparably, in all three neuronal subtypes, the proportion of Twinkle+ dendritic mitochondria are significantly higher (45-62%) than in axons/presynapses.

Taken together, these results demonstrate that most axonal/presynaptic mitochondria lack mtDNA in four different neuronal subtypes of the CNS, whereas in the same neurons, the majority of dendritic mitochondria contain mtDNA.

## A low fraction of axonal mitochondria contain mtDNA as detected by single molecule DNA-FISH

One potential reason for the detection of such a low fraction of nucleoid+ mitochondria in axons of neurons *in vitro* and *in vivo* could be the relatively poor sensitivity of antibody-based detection of mtDNA or cDNA-based expression of mtDNA-associated proteins such as TFAM or Twinkle. We therefore implemented the use of an independent endogenous mtDNA detection method. First, we implemented DNA-fluorescent *in situ* hybridization (DNA-FISH) using probes detecting two mitochondrial DNA encoded genes: Cytochrome b (Cytb) and Cytochrome oxidase 1 (Cox1). DNA-FISH for mtDNA was applied to mature cortical neuron cultures at 21DIV where mito-YFP (matrix-targeted YFP) was sparsely expressed in layer 2/3 CPNs using EUE at E15 (**Fig. 4A-B**). Quantification reveals that only ∼4% of axonal mitochondria are labeled with either Cytb and/or Cox1 probes whereas ∼97% of dendritic mitochondria are positive for both mtDNA probes detecting Cytb and/or Cox1, validating that our DNA-FISH approach can efficiently detect both genes simultaneously (**Fig. 4C-D**). As previously shown *in vitro* and *in vivo* (1, 38, 39), we confirmed that in these culture conditions, dendritic mitochondria of CPNs are large, elongated and fused, whereas axonal mitochondria are small (∼1μm in length) (**Fig. 4E**).

**Figure 4.**
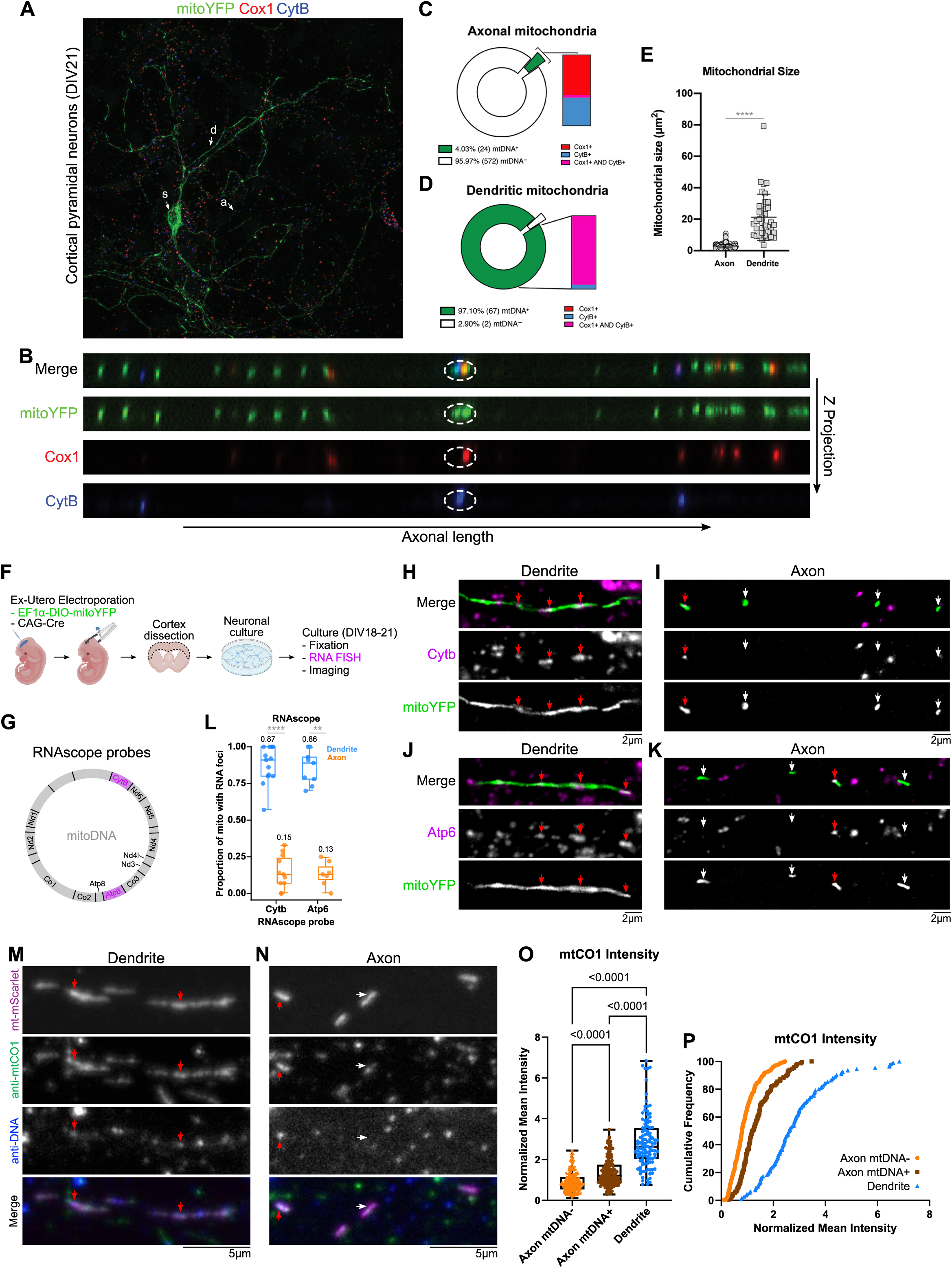
Single molecule DNA-FISH and RNA-FISH reveal that a loew fraction of axonal mitochondria contain either mtDNA or mtDNA-derived mRNA transcripts and lower content of mtDNA-encoded protein mtCO1 in Twinkle- than Twinkle+ axonal mitochondria. (**A-B**) Low magnification (A) of a single cortical pyramidal neuron at 21DIV expressing matrix-targeted YFP (mito-YFP) following DNA-FISH detection of two mitochondria genes Cox1 (red) and Cytochrome B (blue). Abbreviations- s: soma; d: dendrites; a: axon. High magnification (B) confocal z-projection of multiple mitochondria along the axon highlighting a single mitochondrion co-localized with both Cox1 and CytB DNA-FISH signal (circled). (**C-D**) Quantification of Cox1+, CytB+ or double positive mitochondria along the axon (C) and dendrites (D) of CPNs in culture reveals that the majority of axonal mitochondria (>95%) of mitochondria are negative for Cox1 and/or CytB DNA-FISH signals, whereas both mtDNA genes are detected in the majority (97%) of dendritic mitochondria in the same neurons. p<0.0001 (****) by Chi-square test. (**E**) Quantification of mitochondria size (surface area) in axons and dendrites in CPNs imaged for DNA-FISH experiments. Each point is an individual segment. N_axon_ = 596 mitochondria, 24 segments; N_dendrites_ = 60 mitochondria, 6 segments. p<0.0001 (****) by Mann-Whitney test. (**F-G**) Experimental approach (F) used to perform RNA-FISH detection of two mt-DNA encoded mRNAs: *CytB* and *Atp6* (see G) in CPNs sparsely expressing mitoYFP. Briefly, a Cre-dependent plasmid (EF1a-DIO-mitoYFP) was co-electroporated with low amount of a Cre expressing plasmid using EUE at E15, followed by acute dissociation and culture for 18-21DIV. Upon fixation, RNASCOPE^TM^ RNA fluorescent probes were used to detect *Cytb* and *Atp6*. (**H**-**K**) Representative images showing co-localization of *Cytb* (H-I) and *Atp6* (J-K) mRNAs with mitoYFP mitochondria in dendrites (H and J) and axons (I and K) of CPNs *in vitro*. Red arrows indicate mitochondria positive for *Cytb* or *Atp6* mRNAs, whereas white arrows indicated mitochondria negative for the same mRNA probes. (**L**) Quantification of the fraction of axonal and dendritic mitochondria positive for *Cytb* and *Atp6* mRNA fluorescent signal. Each dot represents individual ROIs. Number of mitochondria analyzed: *Cytb* dendrite n=197, *Cytb* axon n=190, *Atp6* dendrite n=129, *Atp6* axon n=162 from three independent cultures. p<0.0001 (****) or p<0.01 (**) by Mann-Whitney test. (**M-N**) Representative images of dendritic (M) and axonal (N) mitochondria in cortical pyramidal neurons maintained for 7DIV and expressing mt-mScarlet (purple) and immunofluorescently stained with antibodies detecting mtCO1 protein (green) and DNA (blue). Orange arrow points to an axonal mitochondrion positive for DNA, while the blue arrow points to an axonal mitochondrion negative for DNA. (**O-P**) Quantification of mean mtCO1 immunofluorescent intensity normalized to the average immunofluorescent intensity of axonal mitochondria negative for DNA. Data displayed as a minimum to maximum box plot with 25^th^, 50^th^ and 75^th^ percentiles marked, colored points represent single mitochondria in each compartment. Statistical analysis: Kruskal-Wallis test. Sample size: mtDNA- axonal mitochondria n=308, mtDNA+ axonal mitochondria n=151, dendritic mitochondria n=110 from three independent cultures.

## The majority of axonal mitochondria lack mtDNA-derived mRNA transcripts and proteins in cortical pyramidal neurons

One testable prediction of the disparity in mtDNA content among axonal mitochondria is that the presence or absence of mtDNA should correlate with the presence or absence of mtDNA-derived mRNA transcripts. To test this, we performed single molecule RNA-FISH using two separate sets of RNA fluorescent probes to detect two mtDNA-encoded mRNAs: *Cytb* and *Atp6* (**Fig. 4G**). Detection of these probes was done in 18-21DIV cortical neuron cultures expressing mito-YFP (introduced with EUE) to visualize dendritic and axonal mitochondria (**Fig. 4F**). In line with our DNA-FISH results, only a small fraction of axonal mitochondria (13-15%) showed positive staining for either RNA probe, while most dendritic mitochondria (>85%) contain several *Cytb* or *Atp6* RNA foci (**Fig. 4H-L**). Importantly, detection of mitochondrial RNA using RNA-FISH was not affected by DNAase treatment (**Fig. S7B**) demonstrating that the RNA-FISH signal we detected are not contaminated by cross-hybridization with mtDNA.

A second testable prediction of the low fraction of axonal mitochondria containing mtDNA is that the abundance of mtDNA-encoded proteins should be lower in mtDNA- axonal mitochondria than mtDNA+ axonal mitochondria. To test this hypothesis, we performed immunocytochemistry with an antibody against mitochondrial encoded Cytochrome c Oxidase subunit 1 (mtCO1) (**Fig. 4M-P**). In DIV7 cultured cortical pyramidal neurons, we observed (1) an ∼3 fold (2.86±0.12) higher fluorescent intensity of mtCO1 staining in dendritic mitochondria compared to axonal mitochondria, and (2) ∼40% (1.38±0.06) increase in mtCO1 immunofluorescence in mtDNA+ axonal mitochondria over mtDNA- axonal mitochondria.

## A low fraction of axonal mitochondria contains mtDNA using quantitative PCR detection from single mitochondria isolated by Scanning Ion Conductance Microscopy (SICM)

Next, we developed an independent way to detect mtDNA in single mitochondria that does not rely on imaging, by modifying and implementing Scanning Ion Conductance Microscopy (SICM) (40). Briefly, single mitochondria were extracted via an epifluorescence microscope equipped with a nanopipette that contains an internal Ag/AgCl electrode inside for monitoring the ionic current between the electrode and cell surface. Femto- to picoliters of liquid can be captured into the pipette by transiently increasing the potential on the electrode (**Fig. 5A**) (41). We isolated individual mitochondria using SICM in CPNs expressing Twinkle-mRuby3 and maintained in culture with a physiological range of glucose concentrations (5 - 10 mM) for 7-9DIV. A mitochondrial matrix-targeted YFP (mito-YFP) was expressed to visualize all the mitochondria in the culture. This approach allows rapid, visually-guided, capture of individual Twinkle+ or Twinkle- mitochondria in axons or dendrites (**Fig. 5B-D, Supplementary Movie 1**). Upon isolation of single axonal or dendritic mitochondria, we performed calibrated quantitative PCR using dual-labeled probes to measure the presence of mtDNA in a highly specific and efficient manner (**Fig. 5E-F**). This analysis confirms that only a small (∼10%) fraction of axonal mitochondria contains qPCR-detectable mtDNA, whereas most dendritic mitochondria contain multiple (up to 10) mtDNA copies (**Fig. 5G-H**).

**Figure 5.**
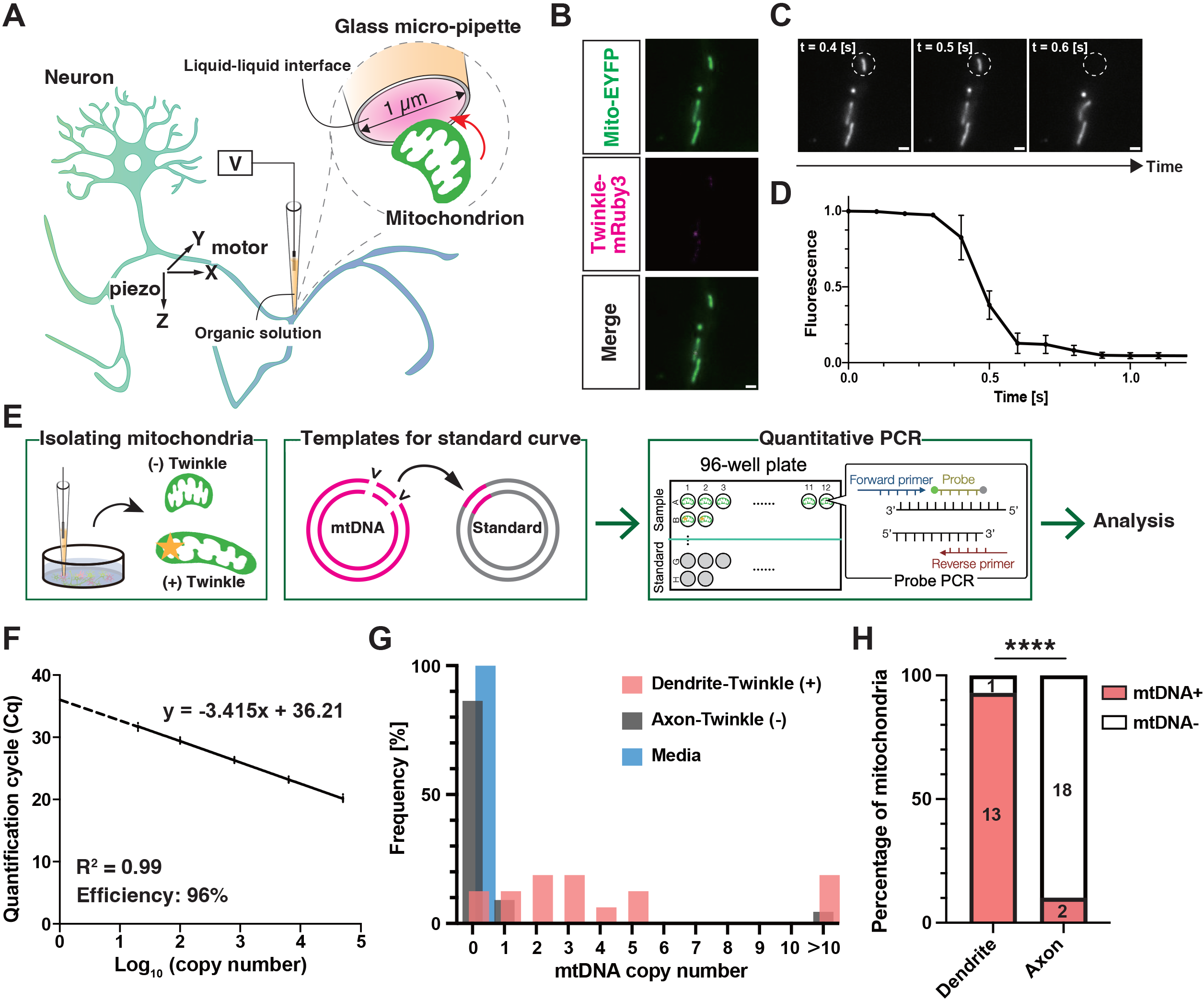
Single axonal mitochondrion isolation coupled with quantitative PCR confirm the low fraction of mtDNA+ mitochondria in axons of CPNs. (**A**) Schematic illustration of single mitochondrion isolation from mouse primary neocortical neurons using a nanopipette with an Ag/AgCl electrode (Scanning Ion Conductance Microscopy; SICM). Upon changing the voltage across the organic solution/aqueous solution interface, an extraction force is applied to a single mitochondrion in 7-9DIV neurons. (**B**) Virtually all neurons were infected with a lentivirus encoding mitoYFP (green), while a fraction of neurons was electroporated with a plasmid encoding Twinkle - mRuby3 (magenta). (**C**) A nanopipette was positioned close to a labeled axonal mitochondrion (dashed circle; t=0.4 s). By changing the voltage, single mitochondrion was trapped into the tip (t=0.5 s) and subsequently removed from the neuron (t=0.6 s). Scale bar, 5 µm. (**D**) Changes of the EYFP fluorescence in areas corresponding to targeted mitochondria were plotted with mean±SEM (n = 7 independent measurements) revealing rapid (<0.3s) isolation of single mitochondrion. (**E-G**) Single Twinkle+ or Twinkle- mitochondria isolated by SICM were collected in individual wells of 96-well plates (E) to perform quantitative PCR calibrated (F) using serially diluted plasmids containing a mtDNA sequence corresponding to the sequence used to determine mtDNA copy number in each isolated axonal mitochondrion by qPCR. This approach allows estimation of mtDNA copy number in all mitochondria isolated from axons or dendrites (G). (**H**) Percentages of mitochondria in which mtDNA were detected (numbers of mitochondria in each category indicated in the bar graphs). **** p < 0.001 by Chi-square test.

## Axonal presynaptic mitochondria lacking mtDNA contain cristae and are not associated with autophagosomes *in vitro* and *in vivo*

Because of their small size (∼1μm in length) and their frequent lack of mtDNA, we next examined the ultrastructural features of axonal mitochondria to determine if they possess cristae, the folded inner mitochondria membrane (IMM) where protein complexes forming the electron transport chain (ETC) are located. To achieve this, we implemented a correlated light-electron microscopy (CLEM) pipeline and determined the ultrastructural features of axonal mitochondria expressing Twinkle or not (**Fig. S5A-B and Supplementary Movie S2**). Three-dimensional (3D) serial EM microscopy reconstructions demonstrate that both fluorescently labeled Twinkle+ and Twinkle- axonal mitochondria have ultrastructurally identifiable cristae in CPNs *in vitro* (**Fig. S5C-D and Supplementary Movie S3**). We also found the clear presence of cristae in presynaptic mitochondria of CPNs *in vivo*, using publicly accessible serial EM datasets (42) (**Fig. S6A and Supplementary Movie S4**). We then confirmed that small axonal mitochondria are rarely (∼15%) associated with autophagolysosomes (mitophagy structures) by visualizing mitochondria, LAMP1 and LC3 simultaneously in 14 DIV cortical axons following EUE at E15 with plasmids encoding mito-mTAGBFP2, Lamp1-mEmerald and LC3-mRuby (**Fig. S6B-C**). Together, these results demonstrate that small axonal and presynaptic mitochondria are distinct, intact organelles.

## Most axonal mitochondria entering the axon and dendrites from the soma lack mtDNA

What are the cellular mechanisms leading to such a low fraction of axonal mitochondria containing mtDNA? We envisioned two potential mechanisms: (1) Recent work demonstrated that in mammalian cell lines, mtDNA replication is coupled with mitochondrial fission to ensure that both ‘daughter’ mitochondria both contain mtDNA (43–45). Therefore, we hypothesized that the high levels of mitochondrial fission factor (Mff)-dependent fission characterizing mitochondria of CPNs axons (1) might be non-replicative, progressively diluting the fraction of mitochondria containing mtDNA with each round of fission and/or (2) a large fraction of mitochondria entering the axon from the soma could already be lacking mtDNA.

We tested the first model, i.e. if a high rate of fission is required for the lack of mtDNA+ mitochondria in axons of CPNs, by blocking mitochondrial fission using shRNA-mediated knockdown of Mff or Fission 1 (Fis1) (1). Mff and Fis1 were chosen as recent work demonstrated that in cell lines, mitochondria can undergo two types of fission events: Mff mediated midzone fission to generate symmetric fission, and Fis1 mediated endzone clipping which occur at the end of elongated mitochondria to generate asymmetric daughter mitochondria (45). We therefore performed *ex utero* electroporation (EUE) at E15.5 with plasmids encoding matrix targeted mScarlet (mt-mScarlet) and either a control shRNA or the previously validated shRNA against either Mff (1) or Fis1 (**Fig. S8**) followed by *in vitro* dissociated culture. At 21DIV, sparsely electroporated CPNs were fixed and immunolabeled for mScarlet and endogenous DNA to visualize mtDNA+ axonal mitochondria (**Figure 6A-C**). We observed no significant change in the percentage of axonal mitochondria containing mtDNA upon either Mff (despite drastic increase in axonal mitochondria size) or Fis1 knockdown, arguing that mitochondrial fission in not a major contributor to the lack of mtDNA in axonal mitochondria of CPNs (**Figure 6D**).

**Figure 6.**
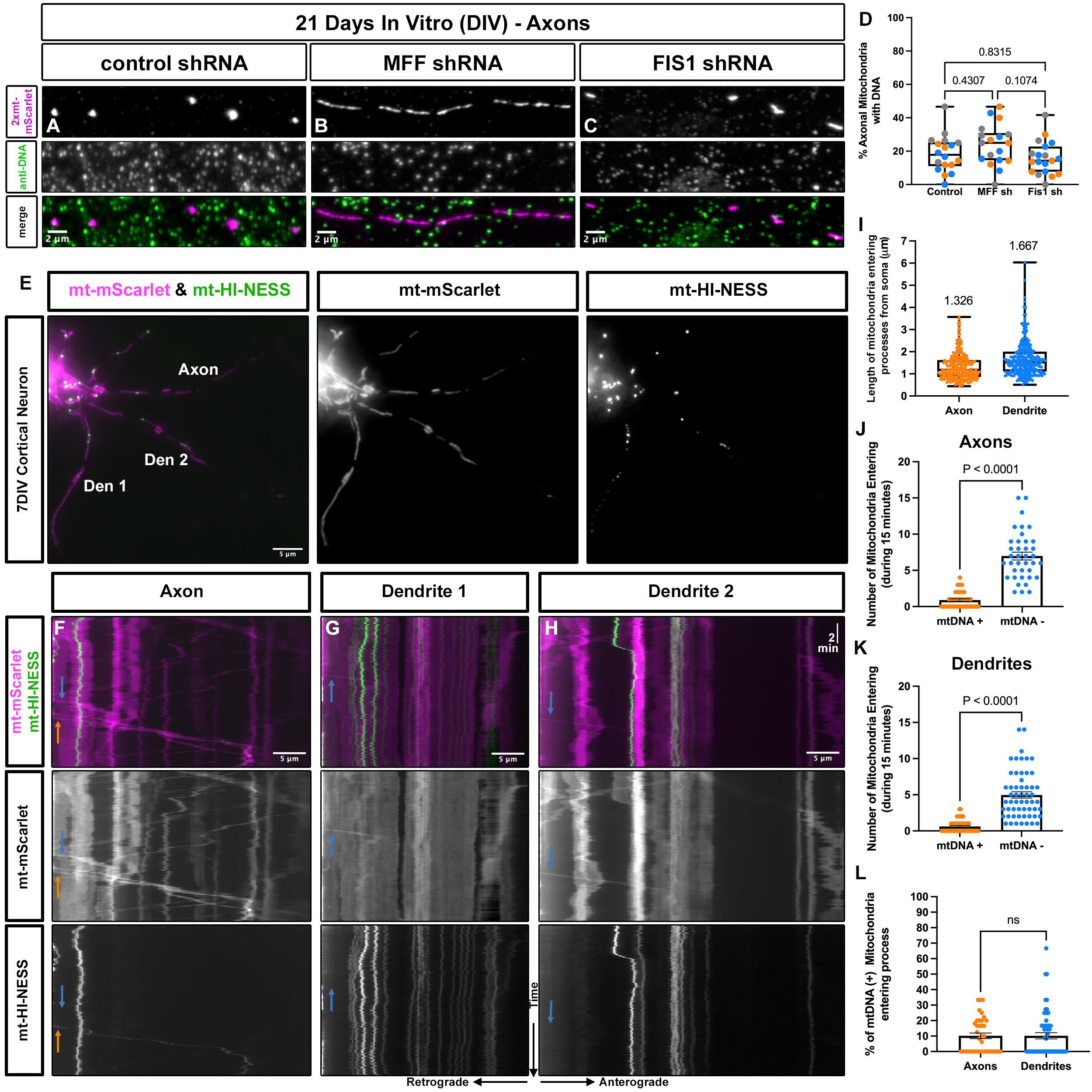
The majority of mitochondria entering axon and dendrites from the soma are small and already lack mtDNA. (**A-C**) Axonal mitochondria in CPNs electroporated with control shRNA are punctate and mostly lack anti-DNA immunofluorescent staining (A). shRNA-mediated knockdown of mitochondrial fission factor (*Mff*) (B) or *Fis1* (C) to block the two main types of fission events does not impact the fraction of axonal mitochondria containing mtDNA arguing that the coupling of mitochondrial fission to mtDNA replication is not a major mechanism for regulating mtDNA levels in the axon. (**D**) Quantification of the percentage of axonal mitochondria containing DNA following knockdown with the indicated constructs. N_control shRNA_ = 335 mitochondria from 18 axon segments from 3 independent cultures. N_Mff shRNA_ = 314 mitochondria from 18 axon segments from 3 independent cultures. N_Fis1 shRNA_ = 362 mitochondria from 19 axon segments from 3 independent cultures. Graphs are minimum to maximum box plots with 25^th^, 50^th^, and 75^th^ percentiles marked, colored points represent data from the single neuronal cultures. Statistical analysis performed with Brown-Forsythe and Welch ANOVA test. (**E**) Representative image of a cortical pyramidal neuron expressing mt-mScarlet (purple) and mt-HI-NESS, a genetically-encoded probe detecting mtDNA in live cells (green) at 7DIV. (**F-H**) Kymographs (time on Y-axis and distance along processes on X-axis) showing trafficking of individual mitochondria and mtDNA labeled with mt-HI-NESS entering the axon (F) or dendrite 1 (G) or dendrite 2 (H) shown in panel E. Orange arrow in F indicates a mt-HI-NESS positive mitochondria whereas blue arrows in F-G indicate mt-HI-NESS negative mitochondria. (**I**) Quantification of the length of mitochondria entering the proximal portion of axons or dendrites. Mean length (microns) is listed above each box plot. (**J**) Quantification of the number of mtDNA+ or mtDNA- mitochondria entering axons during each 15min timelapse sessions. (**K**) Quantification of the number of mtDNA+ or mtDNA- mitochondria entering dendrites during each 15min timelapse sessions. (**L**) Quantification of the fraction of mtDNA+ mitochondria entering axonal and dendritic processes demonstrating that only ∼10% of mitochondria entering proximal portion of either axons or dendrites are positive for mtDNA. p values are shown in the figure using Wilcoxon test (J-K), Mann-Whitney test for L. For panels I-L, n=299 mitochondria entering axons (n=38) from soma, and n=312 mitochondria entering dendrites (n=56) from soma from three independent neuronal cultures.

To test the second model, i.e. to determine whether axonal mitochondria enter the axon already devoid of a nucleoid/mtDNA, we performed timelapse imaging of axonal mitochondria in CPNs expressing matrix-targeted mScarlet (mt-mScarlet) and a recently developed genetically-encoded probe for visualizing mtDNA called mt-HI-NESS which labels mtDNA with the photoconvertible green fluorescent protein Kaede (46). This probe encodes a matrix-targeted (Cox8 mitochondrial targeting sequence), bacterial H-NS DNA binding domain and has been shown to bind to mtDNA. We validated that this mt-HI-NESS reporter colocalizes with DNA staining in both axonal and dendritic mitochondria of cortical PNs maintained *in vitro* (**Fig. S9A-C**). We performed timelapse analysis of mt-HI-NESS-positive and negative mitochondria emerging from the soma entering the proximal portions of dendritic and axonal processes (**Fig. 6E-H**). Interestingly, most mitochondria entering both axons and dendrites from the soma exhibit small size (∼1.3-1.7 μm long; **Fig. 6I**). Furthermore, only a minority (∼10%) of mitochondria entering axonal and dendritic segments from the soma are positive for mt-HI-NESS (**Fig. 6J-L**).

As an alternative approach to label mtDNA, we also performed timelapse imaging of axonal mitochondria in CPNs maintained in culture for 10DIV expressing OMM targeted-mCherry (mCherry-ActA) and Twinkle-Venus to label mtDNA-associated nucleoids (**Fig. S9C-D**). Following 15 minutes of live imaging, we observed between 1 to 8 mitochondria entering the axon from the cell body (**Fig. S9E**). Strikingly, in ∼75% (13/17) of the axons imaged, none of the axonal mitochondria entering the axon during the 15 minutes period of imaging were positive for Twinkle-Venus (**Fig. S9F**). Of the 17 axons imaged, we observed only 5.6% (4/71) of axonal mitochondria entering the axon to be Twinkle-Venus+ (**Fig. S9E**). These results strongly suggest that the majority of biogenesis/fission events operating in the soma to generate small (∼1μm long) mitochondria entering the axon or dendritic processes lack mtDNA.

## F_1_F_0_-ATP synthase (Complex V) in axonal mitochondria functions in reverse-mode extruding H^+^ and consuming ATP to maintain mitochondrial membrane potential

Taken together, our results identify a fundamental difference between axonal and dendritic mitochondria regarding mtDNA content: in all four neuronal subtypes examined above, dendritic mitochondria form a dense, fused, elongated network containing numerous mtDNA+ nucleoids whereas most axonal mitochondria are small (∼1μm in length) and do not contain mtDNA. We previously demonstrated that the striking degree of compartmentalization of mitochondrial structure between axons and dendrites of CPNs observed *in vivo* (2) is controlled by a significant difference in the fusion/fission balance, with Mff-dependent fission being much more prevalent in axons (1). However, our results regarding the dramatic difference in mtDNA content between axonal and dendritic mitochondria raise an important question: are these structural differences reflecting a functional divergence? In eukaryotic cells, the 16.5 kB long mtDNA contains 37 genes, 13 of which encode proteins, 22 encode tRNAs and 2 encode ribosomal RNAs used for mitochondrial translation. The 13 protein-coding genes contained in the mitochondrial genome encode several key proteins within the large complexes composing the electron transport chain such as the ND1-6 subunits of Complex I (NADH dehydrogenase), Cytochrome b (Cytb; Complex III) and Cytochrome c oxidase (Complex IV) and two subunits of the F_1_F_0_-ATP synthase (Complex V). Therefore, one would hypothesize that axonal mitochondria lacking mtDNA would have a less effective ETC and therefore relatively low capacity for oxidative phosphorylation and ATP synthesis.

However, with most axonal mitochondria lacking mtDNA, and lower expression of mtDNA mRNA components encoding oxidative phosphorylation complexes mediating ETC, H^+^ extrusion across the inner mitochondrial membrane (IMM) could be significantly reduced in axonal mitochondria. To determine whether axonal and dendritic mitochondria have distinct functional properties, we first tested whether mitochondrial matrix pH, i.e. H^+^ concentration dynamics, differs in mitochondria found in these two compartments. Thus, we performed EUE at E15.5 to express a plasmid encoding a matrix targeted, fluorescent pH reporter (mt-SypHer; (47)) and a non-pH sensitive matrix targeted HA-mCherry (mt-HAmCherry) to assess basal matrix pH in axonal and dendritic mitochondria of layer 2/3 CPNs maintained in culture for 21DIV. Using live confocal microscopy, single dendritic and axonal segments were imaged from the same CPN allowing paired comparison of the mt-SypHer/mCherry ratio of axonal and dendritic mitochondria (**Fig. 7A**). Both at the individual mitochondrion level (**Fig. 7B**), and when averaged throughout a segment (**Fig. 7C**), axonal mitochondria consistently display a significantly higher SypHer/mCherry ratio, indicative of a more basic pH (lower H^+^ concentration) in the matrix of axonal mitochondria, compared to dendritic mitochondria. To determine whether this more basic matrix pH is the result of altered H^+^ flux by the ETC of axonal mitochondria, we treated CPN cultures expressing mt-SypHer and labeled with a membrane potential dye, tetramethylrhodamine (TMRM; 20 nM loading, 5 nM during imaging), with either a complex III (Antimycin A – 1.25 µM) or complex V (Oligomycin - 1.25 µM) inhibitor and measured variations in axonal mitochondria matrix pH and membrane potential (**Fig. 7D**). As expected with complex III inhibition, the mitochondrial matrix acidified significantly (decreased fluorescence of mt-SypHer, i.e. increased [H^+^] in mitochondrial matrix; (47)) and mitochondrial membrane potential (TMRM) drops rapidly (**Fig. 7E-H**). This suggests that Complex III is functioning, at least to some extent, in axonal mitochondria, contributing to some H^+^ extrusion outside the matrix. However, with Oligomycin treatment, which blocks Complex V (F_1_F_0_-ATP synthase), instead of the expected de-acidification (reduction in matrix [H^+^]) and hyperpolarization of the membrane potential, we observed again that the matrix acidifies and membrane potential drops (**Fig. 7E-H**). This strongly argues that in axonal mitochondria, the F_1_F_0_-ATPase is working in reverse mode, extruding H^+^ out of the matrix, and that axonal mitochondria hydrolyze ATP at steady state, thereby participating in the maintenance of mitochondrial membrane potential.

**Figure 7.**
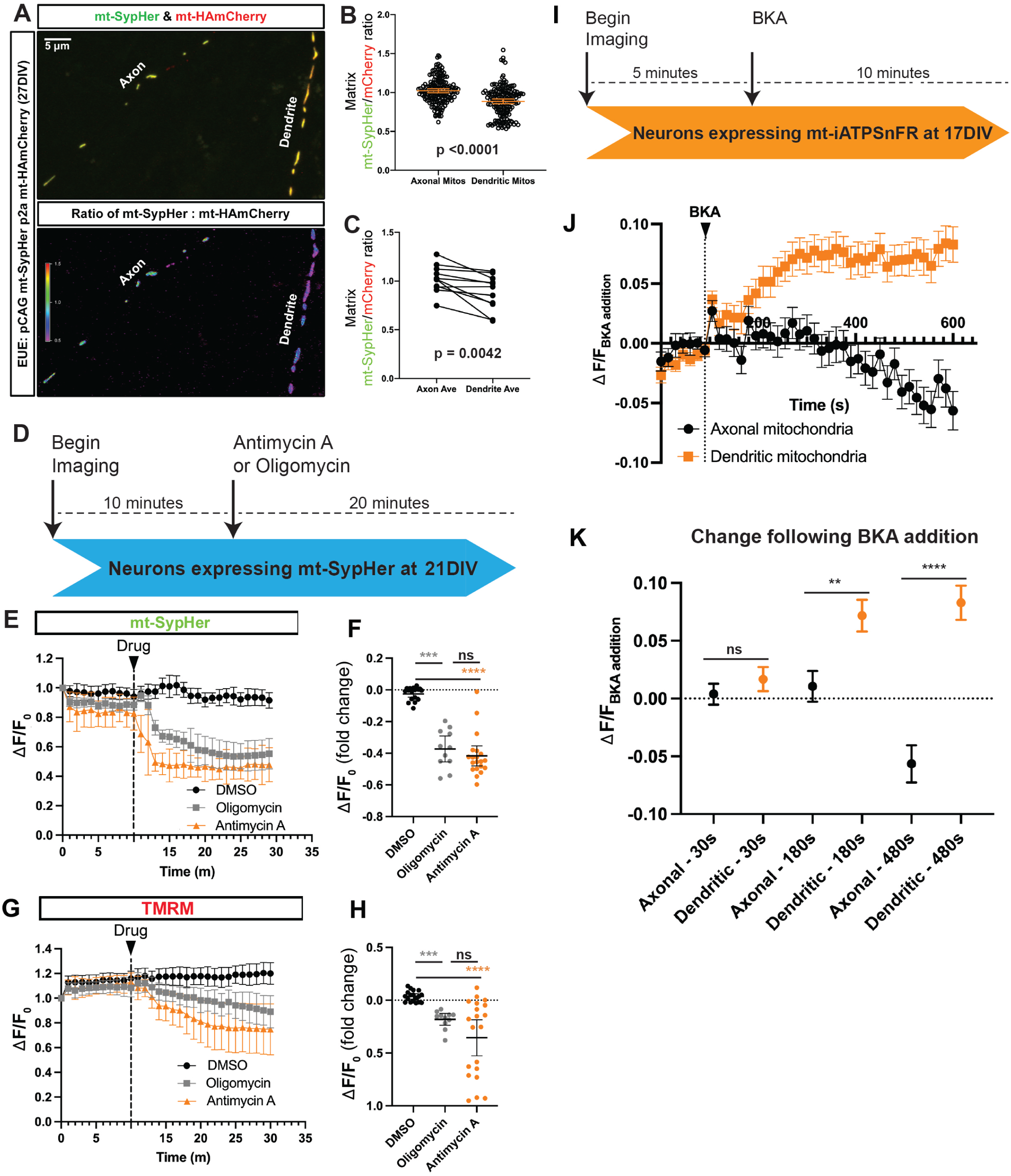
Axonal mitochondria in CPNs run Complex V in reverse and consume ATP to maintain membrane potential. (**A**) Axonal and dendritic mitochondria from the same neuron expressing mitochondrial matrix targeted SypHer (mt-SypHer) and HAmCherry (mt-HAmCherry) in culture for 21DIV. (**B-C**) Quantification of the SypHer to HA-mCherry ratio for individual, pooled mitochondria (B) or of paired axon and dendrite averages (C). A significant increased mt-SypHer/mCherry ratio in axonal compared to dendritic mitochondria suggests that the matrix of axonal mitochondria is more basic (lower H^+^ concentration) than the matrix of dendritic mitochondria (**D**) Scheme for measuring axonal mitochondria responses following addition of either Antimycin A or Oligomycin. (**E-F**) Quantification of mt-SypHer fluorescence over time before and following addition of Antimycin A or Oligomycin (E) and average fold change of mt-SypHer fluorescence in individual axons (F). (**G-H**) Quantification of TMRM fluorescence over time before and following addition of either Antimycin A or Oligomycin (G) and average fold change of TMRM fluorescence in individual axons (H). Both Antimycin A and Oligomycin elicit acidification and loss of membrane potential in axonal mitochondria. (**I**) Scheme for assaying mitochondrial matrix ATP levels using matrix-targeted iATPSnFR1.0 (mt-iATPSnFR) in axonal and dendritic mitochondria of CPNs in culture at 17DIV. (**J**) Quantification of the change in matrix-targeted mt-iATPSnFR fluorescence in either dendritic mitochondria (orange) or axonal mitochondria (black) before and following BKA addition showing that dendritic matrix ATP increases with ANT blockade while axonal matrix ATP decreases with ANT blockade. (**K**) Fold change of mt-iATPSnFR 30 s, 180 s or 480 s after BKA addition in matrix of axonal and dendritic mitochondria. Scale bar in A is 5 μm. Error bars are SEM. **p≤ 0.01, *** p≤ 0.001, **** p≤ 0.0001. Kruskal-Wallis test for B, D, E and G. Paired t test for C. N_mt-SypHer axonal_ = 167 mitochondria from 11 neurons, 3 independent cultures, N_mt-SypHer dendritic_ = 162 mitochondria from 11 neurons, 3 independent cultures for B-C; N_DMSO_ = 18 axons from 3 independent cultures, N_Antimycin A_= 19 axons from 3 independent cultures, N_Oligomycin_= 11 axons from 3 independent cultures for E-H; N_BKA axonal_ = 129 mitochondria from 13 neurons, 3 independent cultures, N_BKA dendritic_= 110 mitochondria from 13 neurons, 3 independent cultures for J-K.

The capacity of the F_1_F_0_-ATP synthase to function in a reverse way, promoting ATP hydrolysis to extrude H^+^ out of the matrix is limited by the expression of ATP5IF1 (also called ATPIF1) (48). We reasoned that ATP5IF1 could be expressed at lower levels in axonal than in dendritic mitochondria to enable the F_1_F_0_-ATP synthase to function in a reverse way. To test this, we performed immunofluorescent detection of ATP5IF1 in optically isolated axonal and dendritic mitochondria, labeled with mito-YFP, in cortical PNs *in vitro* (**Fig. S10A-C**). Our results demonstrate a significantly lower abundance of ATP5IF1 protein in axonal compared to dendritic mitochondria (**Fig. S10D-F**).

To test more directly if the F_1_F_0_-ATPase of axonal mitochondria of CPNs function in reverse at steady state, we measured mitochondrial matrix ATP dynamics following inhibition of the adenine nucleotide translocase (ANT) with Bongkrekic acid (BKA; 50 µM) (49). We expressed a matrix targeted iATPSnFR1.0 (mt-iATPSnFR;(50)) fused with mScarlet using EUE at E15.5, then at 17DIV, we performed live imaging before and after BKA application (**Fig. 7I**). Since iATPSnFR is pH sensitive, we first confirmed that over the time frame of the imaging experiment matrix pH is not significantly altered by BKA addition by imaging mitochondria located in the soma (**Fig. S11**). We then imaged mt-iATPSnFR dynamics in both dendritic and axonal mitochondria before and following addition of BKA and observed an increase in the mt-iATPSnFR signal of dendritic mitochondria, indicative of ATP accumulation in the matrix following ANT inhibition (**Fig. 7J-K**). In axonal mitochondria, mt-iATPSnFR instead decreased over time, suggesting that ATP is consumed in the matrix of axonal mitochondria at resting state (**Fig. 7J-K**).

## DISCUSSION

Mitochondria were recently shown to display distinct patterns of dynamics and morphology in dendrites and axons (1, 27), but the functional correlates of these structural observations, if any, at the level of neuronal metabolism remained unknown. In the present study, we show that axonal mitochondria not only display uniformly small size but also that the majority of these axonal mitochondria lack mtDNA/nucleoids, have low levels of mtDNA-encoded mRNAs (*Cytb* and *Atp6*) and protein (mtCO1) content, low levels of ATP5IF1 and hydrolyze ATP instead of promoting oxidative phosphorylation. Taken together with previously published results demonstrating the profound differences in mitochondria structure between axons and dendrites in mammalian long-projecting cortical pyramidal neurons (1, 2, 51), our results demonstrate that these striking structural differences have a functional correlate. In dendrites of CPNs, mitochondria are elongated and highly fused, with their network occupying a large fraction of the dendritic arbor and are highly metabolically active generating ATP while containing numerous mtDNA-nucleoids. In contrast, axonal mitochondria in the same neurons are small (∼1μm long), mostly lacking mtDNA, and hydrolyze ATP through the reverse action of F_1_F_0_-ATP synthase (Complex V) to maintain a physiological hyperpolarized membrane potential range. One potential explanation for axonal mitochondria of CPNs using Complex V to consume ATP and protrude H^+^ outside the matrix at steady state may be that some of the key functions of presynaptic axonal mitochondria, such as MCU-dependent Ca^2+^ import, require mitochondrial membrane potential to be maintained within a physiological, hyperpolarized range, despite having suboptimal ETC function (52).

The surprising model emerging from our results is that the main source of ATP in axons of mammalian CPNs could be intrinsic glycolysis (53), whereas mitochondria in the dendritic compartment are fully competent to generate ATP through oxidative phosphorylation. We propose that the axonal/presynaptic mitochondria identified in this study that lack mtDNA and hydrolyze ATP represent a new class of specialized mitochondria as recently described in other cell types or sub-cellular compartments for their specialized structural and/or functional properties (54–56).

What could be the biological adaptation of such a drastic difference between the mtDNA content and ATP generation capacity between axonal and dendritic mitochondria in mammalian CPNs? We speculate four potential scenarios to explain this drastic level of compartmentalization of mtDNA content and mitochondrial function between dendrites and axons of CNS neurons: (1) the cytoplasmic volume at individual presynaptic bouton is extremely small (in the order of hundreds of cubic nanometers) and therefore, in such extremely small volumes, the accumulation of mitochondria-derived reactive-oxygen species (ROS), a byproduct of highly functional ETC, could be damaging to protein complexes involved in neurotransmission; (2) mitochondria with highly functioning electron transport chain and high levels of oxidative phosphorylation have been proposed to reach temperatures approaching 50°C (57, 58). Since axonal mitochondria are located just a few hundreds of nanometers from the active zone at presynaptic boutons, where SNARE-mediated presynaptic vesicles are docked near the plasma membrane, and SNARE-mediated exocytosis is highly temperature-dependent (59), we speculate that the axonal/presynaptic mitochondria lacking mtDNA and hydrolyzing ATP might not generate high temperatures and therefore would not interfere with SNARE-mediated presynaptic vesicle fusion; (3) Axonal mitochondria are more inclined towards electron transport chain-independent anabolic functions, such as amino acid biosynthesis (60, 61) including glutamate generation, the primary excitatory neurotransmitter used by CPNs and/or to fuel local protein synthesis which is prevalent, and often associated with mitochondria, not only in dendrites (11, 17) but also in distal portions of the axon in neurons (62, 63); (4) recent work suggests that mtDNA-mediated activation of the cGAS-STING pathway triggers low grade inflammation and senescence-associated secretory phenotypes found in aging and neurodegeneration (64, 65). Therefore, limiting the abundance of mtDNA along CNS axons might have been selected during evolution to limit cGAS-STING engagement during the lifetime of long-lived vertebrates in a neuronal compartment where mitophagy is prevalent (66, 67). Future investigations will need to test these hypotheses

Throughout this study, we employed four independent approaches to estimate the fraction of mitochondria containing mtDNA in axons and dendrites of four distinct neuronal subtypes within the mouse CNS. Our results provide consistent estimates in cortical pyramidal neurons both *in vitro* and *in vivo* as well as PV+ and SST+ cortical inhibitory interneurons and dopaminergic neurons located in SNc *in vivo*. However, each of these four independent techniques have different sensitivity for mtDNA detection and therefore provide a range of values for the fraction of axonal mitochondria containing mtDNA: (1) using single-photon confocal microscopy to detect DNA and/or Twinkle labeling in **Fig. 1C** and **Fig. 3**, we obtained 12.6% for CPNs at 15DIV cultured with 25mM glucose and 13.2% for CPNs at 18DIV cultured with 2.5mM glucose *in vitro*, 24.6% for layer 2/3 PNs *in vivo* using Twinkle (**Fig. 1F**) as well as 16.5% from the *in vivo* TFAM labeling experiment (**Fig. S2**) and 21.4% and 13.8% in axons of cortical PNs *in vitro* based on Twinkle+ mitochondria at presynaptic (vGlut1+) or outside presynaptic (vGlut1-negative) boutons respectively (**Fig. S2G**); using either fluorescently-tagged Twinkle expression or DNA immunolabeling we obtained 29-30% in PV-INs, 15-20% in SST-INs and 8-12% in dopaminergic neurons of the SNc (2) using STED 2-color microscopy detecting mtDNA in axons of rat hippocampal neurons at 14-16DIV *in vitro* we obtained 24.7% (**Fig. 3F**); (3) 4% using DNA-FISH in mouse cortical PNs *in vitro* (**Fig. 4C**); (4) finally, our scanning ion conductance microscopy (SICM) to collect individual axonal Twinkle-negative mitochondria coupled with quantitative PCR for mtDNA was used as a highly sensitive and represents a stringent way to estimate the false discovery rate (FDR) linked to using Twinkle-negative mitochondria as an estimate of the absence of mtDNA (**Fig. 6**). This result demonstrate that this FDR is only ∼10%, i.e. only 10% of axonal mitochondria negative for Twinkle contain one copy of mtDNA, therefore providing strong confidence that the lack of fluorescently tagged Twinkle is a reliable marker for the absence of mtDNA.

Therefore, both *in vitro* and *in vivo*, the estimated range of mtDNA-negative mitochondria in axons is 4-25%, which justifies the statement that, in the four CNS neuronal subtypes examined in this study, the majority (75-96%) of axonal mitochondria lack mtDNA. Importantly, using the same four independent approaches in the same four neuronal subtypes, we report the inverse results: the fraction of dendritic mitochondria containing mtDNA and in fact multiple nucleoids, ranges from 60-95%, with a high content of mtDNA-encoded mRNAs and protein, and high OXPHOS activity.

A recent preprint reported using mostly mass spectrometry-based proteomic approaches that axonal mitochondria are significantly ‘impoverished’ for OxPhos encoding proteins compared to dendritic mitochondria in mouse CNS neurons (68). Future studies will need to extend these results to other neuronal subtypes of the CNS and in other species, including both vertebrate and invertebrates and, critically (1) identify the molecular mechanisms leading to this striking degree of compartmentalization of mtDNA content and functional properties of mitochondria in ATP generation between axons and dendrites and (2) test if this is also observed for neurons located in the peripheral nervous system (PNS), such as sensory or motor neurons that extend axons outside the nervous system, where mitochondrial trafficking and functional/metabolic requirement might be different than in CNS neurons.

## Material and methods

### Mice

All animals were handled according to protocols approved by the Institutional Animal Care and Use Committee (IACUC) at Columbia University, the Oklahoma Medical Research Foundation, University of Tokyo. Time-pregnant CD1 or ICR females were purchased from Charles Rivers. At the time of *in utero*, or *ex utero* electroporation (E15.5), littermates were randomly assigned to experimental groups without regard to their sex. DAT-Cre mice (B6.SJL-*Slc6a3^tm1.1(cre)Bkmn^*/J; Cat# 006660), PV-Cre mice (B6.129P2-*Pvalb^tm1(cre)Arbr^*/J;Cat# 017320) and SST-Cre mice (*Sst^tm2.1(cre)Zjh^*/J; Cat# 013044) were purchased from Jackson Laboratory.

### Lentivirus production

HEK293T cells (RIKEN BRC, RCB2202) were co-transfected with shuttle vectors, VSV-G and Δ8.9 using FuGENE transfection reagent (Promega, E2311). 24 hours after transfection, the media was exchanged with fresh Neurobasal media (Gibco, 21103049), and 48 hours later, supernatants were harvested, spun at 2500×g to remove debris and filtered through a 0.45 μm filter (Membrane Solutions, PES025045). The filtered supernatant was concentrated ×20 using an Amicon® Ultra-15 centrifugal filter device (molecular weight cut-off 100 kDa, UFC910024, Merck Millipore Ltd.), which was centrifuged at 4,000×g for 20 minutes at 4°C. The concentrated samples were diluted with 1×PBS (Nissui, 05913) and stored at −80°C.

### Plasmids

FUW mito-EYFP was created from FUIGW (kind gift from Dr. Yukiko Gotoh) by replacing the IG sequence with mito-EYFP sequence using EcoR1 and BsrG1 sites. pCAG HAmCherry-ActA was created by PCR of the DNA encoding HAmCherry and subcloning it 3’ to the CAG promoter but 5’ to the ActA mitochondrial targeting sequence (69, 70). pCAG Twinkle-Venus was created by PCR of the DNA encoding mouse Twinkle from a neuronal mouse cDNA library, and subcloned 3’ to the CAG promoter but 5’ to the DNA encoding Venus YFP via Infusion cloning. pCAG Twinkle-mRuby3 was created from pCAG Twinkle-Venus by replacing the Venus YFP sequence with mRuby3 via infusion cloning, using Age1 and Not1 sites. pCAG NF186-pHluorin was created by PCR of the DNA encoding mouse NF186 from a neuronal mouse cDNA library in two sections an N-terminal and C-terminal segment, which were subcloned 3’ to the CAG promoter but 5’ or 3’ respectively to the DNA encoding pHluorin via Infusion cloning. pCAG mt-SypHer was created by PCRing the DNA encoding mt-SypHer from Addgene plasmid #48251 (a gift from Nicolas Demaurex), and subcloning it 3’ to the CAG promoter. pCAG mt-SypHer p2a mt-HAmCherry was created by subcloning the DNA encoding 2xmt-HAmCherry 3’ to pCAG mt-SypHer via Infusion cloning. pCAG 4xmt-iATPSnFR1.0 was created by PCR of the DNA encoding iATPSnFR1.0 from Addgene Plasmid #102556 (a gift from Baljit Khakh), and subcloning it 3’ to the 4x synthetic mitochondrial targeting sequence in Addgene plasmid #66896 (a gift from Georg Ramm). This whole DNA encoding the 4x-mt iATPSnFR1.0 was then subcloned 3’ to the CAG promoter via infusion cloning. pCAG HA-mFis1 was created via PCR of the DNA encoding mouse *Fis1* from a neuronal mouse cDNA library, and sub-cloned 3’ to the CAG promoter and a HA tag. pLV-Puro-hPGK-mtKaede-Hi-NESS was a generous gift from Dr. Timothy Shutt to Dr. Tommy Lewis.

### *In utero* electroporation

A mix of endotoxin-free plasmid preparation (2 mg/mL total concentration) and 0.5% Fast Green (Sigma) was injected into one lateral hemisphere of E15.5 embryos using a Picospritzer III (Parker). Electroporation (ECM 830, BTX, CUY21EDITⅡ) was performed with gold paddles to target cortical progenitors in E15.5 embryos by placing the anode (positively charged electrode) on the side of DNA injection and the cathode on the other side of the head. Five pulses of 45 V for 50 ms with 500 ms interval were used for electroporation. Animals were sacrificed 21 days after birth (P21) by terminal perfusion of 4% paraformaldehyde (PFA, Electron Microscopy Sciences) followed by overnight post-fixation in 4% PFA.

### *Ex utero* cortical electroporation

A mix of endotoxin-free plasmid preparation (2-5 mg/mL) and 0.5% Fast Green (Sigma) mixture was injected using a Picospritzer III (Parker) into the lateral ventricles of isolated heads of E15.5 mouse embryo, and electroporated using an electroporator (ECM 830, BTX) with four pulses of 20V or five pulses of 22 V (for the single mitochondria isolation experiments) for 100 ms with a 500 ms interval as described previously (71). Following *ex utero* electroporation, we performed dissociated neuronal culture as described below.

### Primary neuronal culture

Embryonic mouse cortices (E15.5) were dissected in Hank’s Balanced Salt Solution (HBSS) supplemented with HEPES (10 mM, pH7.4), and incubated in HBSS containing papain (Worthington; 14 U/ml) and DNase I (100 μg/ml) for 20 min at 37°C. Then, samples were washed with HBSS, and dissociated by pipetting. Cell suspension was plated on poly-D-lysine (1 or 0.2 mg/ml, Sigma)-coated glass bottom dishes (MatTek) or coverslips (BD bioscience) in Neurobasal media (Invitrogen) containing B27 (1x), Glutamax (1x), FBS (2.5%) and penicillin/streptomycin (0.5x, all supplements from Invitrogen). Every 5 to 7 days, one third of the media was exchanged with supplemented Neurobasal media without FBS.

### Primary neuronal culture at a physiological glucose concentration

Mouse neurons were cultured for the first 4 DIV in Neurobasal media containing B27 (1x), Glutamax (1x), FBS (2.5%). At 4DIV, the half of the media was exchanged with a supplemented BrainPhys Imaging Optimized media (STEMCELL Technologies) without FBS. Half-medium changes were performed every 3 to 4 days.

### AAV injections

For results shown in Figure 3, AAV were produced at 6×10^12^ -4×10^13^ vg/ml and introduced in the mouse brain using stereotaxic injections of AAV9-EF1α::DIO-mitoYFP (1:10) and AAV9 hSyn::DIO-Twnk-mRuby (1:5) were administered (350 nl) bilaterally into the Substantia Nigra pars compacta (SNc) of 3 -month-old DAT-Cre knockin mice via stereotaxic injection under isoflurane anesthesia. Infusions (350 nl/hemisphere) were performed at a rate of 140nl/min using a 5 μl Hamilton glass syringe, with the needle remaining in place for 2 min post-injection. The coordinates for bilateral injection relative to Bregma were AP: −2.910 mm, ML: ±1.2 mm, and DV: −4.9 mm. Following viral expression for two weeks, mice were transcardially perfused with 30 mL of 2% PFA/0.075% GA in 1xPBS. Post-fixed brains were sectioned at 120 µm via vibratome (Leica VT1000 S) and incubated in a blocking solution containing 0.2% Triton X-100, 2% BSA, and 5% Normal Goat Serum. Sections were then stained with primary antibodies in blocking solution. Following washing, the sections were incubated in Alexa-conjugated secondary antibodies (1:2000) in block, and then mounted on slides and coverslipped with Aqua PolyMount (Polymount Sciences).

Similarly, delivery of Cre-dependent mitoYFP and Twnk-mRuby AAV9 into the somatosensory cortex of 2-3-month-old PV-Cre or SST-Cre knockin mice was performed via stereotaxic injection under isoflurane anesthesia. A small circumference of ∼2 mm in diameter was opened in the skull of animals, centered to coordinates AP: -1.8 mm and ML: -2.3 mm relative to Bregma. Within the opening, multiple injections of 64 nl of AAV were performed at a rate of 10 nl/sec with a Nanoject^TM^ (#3-000-207, Drummond Scientific Company), at a depth of 0.5 mm from the top of the exposed cortex. After two weeks of expression, animals were intracardially perfused with 15 ml of 4% PFA in 1xPBS. Dissected brains were post-fixed for 24 hs and 100 µm-thick brain slices were obtained using a vibratome. Sections were then incubated with the indicated primary antibodies diluted in PBS-Triton 0.3% containing 10% Normal Goat Serum for 48 hs at 4°C with agitation. Following primary antibody, sections were washed in 1xPBS, incubated with Alexa-conjugated secondary antibodies (diluted 1:500 in the same buffer used for primary antibodies) for 2 hs at RT, washed again in 1xPBS, and mounted using Citifluor mountant solutions (#17977-150 and # AF100-5, Electron Microscopy).

Primary antibodies included: chicken anti-GFP (Aves), rabbit anti-DsRed (Takara Bio), chicken anti-Tyrosine hydroxylase (Abcam), rabbit anti-Parvalbumin (#PV27, Swant), guinea pig anti-Somatostatin (#366004, Synaptic Systems), and mouse anti-DNA (American Research Products, Inc), all 1:2000.

### Immunocytochemistry

Primary culture - Cells were fixed for 10 minutes at room temperature in 4% (w/v) paraformaldehyde (PFA, EMS) in PBS (Sigma), then incubated for 30 minutes in 0.1% Triton X-100 (Sigma), 1% BSA (Sigma), 5% Normal Goat Serum (Invitrogen) in PBS to permeabilize and block nonspecific staining, after washing with PBS. Primary and secondary antibodies were diluted in the buffer described above. Primary antibodies were incubated at room temperature for 1-2 hours and secondary antibodies were incubated for 30 minutes at room temperature. Coverslips were mounted on slides with Fluoromount G (EMS). Primary antibodies used for immunocytochemistry in this study are chicken anti-GFP (5 μg/ml, Aves Lab – recognizes GFP and YFP), mouse anti-HA (1:500, Covance), rabbit anti-RFP (1:1,000, Abcam – recognizes mTagBFP2, DsRED and tdTomato), mouse anti-DNA (1:200, American Research Products Inc). All secondary antibodies were Alexa-conjugated (Invitrogen) and used at a 1:2000 dilution. Nuclear DNA was stained using Hoechst 33258 (1:10,000, Pierce)

ATPIF1 staining – Cells were fixed with ice cold 2%PFA/0.075% GA for 10 minutes on ice, then permeabilized with 0.1% Triton X100 in 1xPBS supplemented with 22mg/mL glycine for 10 minutes. Cells were then blocked with 1xPBS supplemented with 0.1% Tween, 1% BSA and 5% NGS for 1 hour. Mouse anti-ATPIF1 (ThermoFisher, 1:1000) was incubated in block overnight at 4C. Following washing with 1xPBS, the sample was incubated in secondary antibody at 1:1500 for 2 hours at room temperature. mtCO1 staining – Cells were fixed with freshly made 4% PFA for 30 minutes at room temperature. Following washing with 1xPBS, the cells were permeabilized and blocked with 0.2% Triton X100, 2% BSA and 5%NGS in 1xPBS for 1 hour. Mouse IgG2a anti-MTCO1 (Invitrogen 1:200), mouse IgM anti-DNA (American Research Products 1:200), and rabbit anti-DsRED (1:1000 Takara Bio) were incubated in block for 2 hours at room temperature with shaking. Following washing with 1xPBS, the samples were then incubated in block with secondary antibody (Invitrogen) at 1:1000 for 1 hour at room temperature. Brain sections - Post fixed brains were sectioned via vibratome (Leica VT1200) at 100 µm. Floating sections were then incubated for 2 hours in 0.4% Triton X-100, 1% BSA, 5% Normal Goat Serum in PBS to block nonspecific staining. Primary and secondary antibodies were diluted in the buffer described above. Primary and secondary antibodies were incubated at 4°C overnight. Sections were mounted on slides and coversliped with Aqua PolyMount (Polymount Sicences, Inc). Primary and secondary antibodies are the same as above.

### Immunohistochemistry

For the section of mouse cortex analyzed in Figure 1f, mitochondria were labeled by expression of Tom20-Halotag. Post-fixed brains were sectioned via vibratome (Leica VT1200) at 50 µm. Sections were then incubated for 1 hour in 0.3% Triton X-100 (Nacalai), 3% Normal Goat Serum, 100 mM Glycine (Nacalai), and 0.03% NaN_3_ (Nacalai) in PBS. The primary antibody (rat anti-GFP, 1:500, Nacalai) for enhancing signals from Twinkle-Venus and 400 nM Halotag ligand conjugated with Janelia Fluor 549 NHS ester (Tocris) were diluted in the buffer described above and incubated at 4°C overnight. The secondary antibody (Alexa488-conjugated, 1:500, Invitrogen) was diluted in the same buffer with 0.02% DRAQ5 (BSU) and incubated at 4°C overnight. Sections were mounted on slides and mounted with Fluoromount-G (SouthernBiotech).

### Imaging

All fixed samples were imaged on a Nikon Ti-E microscope with an A1 confocal except for Figure S2a-b were we used a Nikon Ti2 Eclipse microscope with a Nikon AX confocal microscopy and a Nikon Spatial Array Confocal (NSPARC) detector. All equipment and solid-state lasers (Coherent, 405 nm, 488 nm, 561 nm, and 647 nm) were controlled via Nikon Elements (NIS) software. Nikon objectives used include 20x (0.75NA), 40x (0.95NA) or 60x oil (1.4NA) or 100x oil (NA 1.45). Optical sectioning was performed at Nyquist for the longest wavelength. Analysis of intensity, mitochondrial length and occupancy were performed in Nikon Elements.

Images analyzed for experiments in Figure 3g-h, 3m-n, and S4e-f were collected on a Zeiss LSM 880 microscope equipped with Airyscan detectors, plan-Apochromat 20x and 63x 1.4-NA oil objectives, and Zeiss ZEN software.

Live timelapse imaging in Figure 2e-l was performed on mouse cortical pyramidal neurons electroporated *ex utero* (EUE) at E15 and imaged at 7-21DIV with EMCCD (Andor, iXon3-897) or sCMOS (Hamamatsu Orca Fusion) on an inverted Nikon Ti-E microscope or Nikon Ti2-E (40x objective NA0.95 with 1.5x digital zoom or 60x objective NA1.4) with Nikon Elements. 488 nm and 561nm lasers shuttered by Acousto-Optic Tunable Filters (AOTF) or 395 nm, 470 nm, and 555 nm Spectra X LED lights (Lumencor) were used for the light source, and a custom quad-band excitation/dichroic/emission cube (based off Chroma, 89400) followed by clean up filters (Chroma, ET435/26, ET525/50, ET600/50) were applied for excitation and emission. We used cHBSS as the imaging solution.

For mt-HI-NESS imaging in Fig. 2e-l, at 6DIV, primary cortical neurons cultured on glass bottom dishes were transfected with the following plasmid vectors: pLV-Puro-hPGK-mito-Kaede-HI-NESS (20 ng), pCAG 2xmt-mScarlet (20ng) and pKSS empty vector (960ng) via Lipofectamine 2000. At 7DIV, individual neurons were live imaged by widefield fluorescence microscopy at 60x magnification every 0.5 or 5 seconds for 15 minutes. Individual mitochondria (2xmt-mScarlet) were tracked upon entry into neuronal processes and were analyzed for mtDNA content (Mt-Kaede-HI-NESS). Image acquisition, processing and analysis was performed in Nikon Elements AR software. Nikon’s proprietary “Denoise.ai” plugin was used to enhance detection of mtDNA puncta.

In Figure 7, Tetramethylrhodamine (TMRM, Sigma) imaging was done on neuronal cultures incubated with 10 nM TMRM for 20 minutes at 37°C to load the cells before imaging started. TMRM was maintained in the imaging buffer at 5 nM throughout the experiment. For experiments using mitochondrial toxins, cells were imaged for a base line period then the indicated drug was bath applied at the indicated time points. Antimycin A and Oligomycin were used at 1.25 µM final concentration. Bongkrekic Acid (BKA) was used at 50 µM final concentration. Images were analyzed using the time series module in NIS Elements. Full-length mitochondria were marked by a freehand selection tool intensities were measured. After intensities were corrected for background subtraction, ΔF values were calculated from (F-F_0_).

### Single mitochondria extraction using Scanning Ion Conductance Microscopy (SICM)

Single mitochondria extraction was performed using Scanning Ion Conductance Microscopy (SICM) (40, 41). Our customized SICM consists in an epifluorescence microscope equipped with a homemade nanopipette system with a 40 × 40 μm travel range XY piezo stage (PK2H100-040U, THK precision) and a homemade 9.1 μm travel range Z piezo stage equipped with a piezoelectric actuator (AE0505D08, NEC Tokin) for controlling the nanopipette along the Z-axis. A stepping motor stage with a travel range of 30 mm (KXG06030-G, SURUGA SEIKI) was used for coarse positioning of the nanopipette along the Z-axis. The XY and Z piezo stages were operated by a capacitive sensor-controlled closed-loop piezo controller (NCM7302C, THK precision) and an open loop piezo driver (PH103, THK precision), respectively. The pipette current was detected via a homemade 1 GΩ feedback resistance current amplifier. The stable power supply (LP5392, NF) was used as a homemade current amplifier. The holding voltage for the ion current measurement was supplied by a Digital Analog converter of a field-programmable gate array (USB-7855R OEM, National Instruments) to the Ag/AgCl electrode placed in a solution around the cell.

The glass nanopipettes (inner radius, 1.0 µm) were fabricated from a borosilicate glass capillary (GC100F-15, Harvard Apparatus) using a CO_2_ laser puller (model P-2000, Sutter Instruments) and were filled with a solution of 1–2 dichlorethane containing 10 mM tetrahexylammonium tetrakis(4-chlorophenyl)borate (THATPBCl). Ag/AgCl electrodes were inserted into the micropipette.

To visualize all the mitochondria in a dish, cells electroporated with Twinkle-mRuby3 were infected with lentivirus carrying Mito-EYFP. Voltage was applied to the liquid-liquid interface between dichlorethane in the nanopipette and the culture media in the dish to control the dichlorethane surface tension. To prevent the solution from flowing into the nanopipette before extraction, the voltage was kept at +0.5 V vs Ag/AgCl. After positioning the nanopipette tip close to a target mitochondrion, the voltage was changed to -1.0 V for 300-500 ms to raise the oil-water interface and extract individual mitochondrion into the nanopipette. After confirming the disappearance of the target mitochondrial signal from the cell, the mitochondrion was collected in a 96-well plate (Greiner, 669285) by breaking off the tip of nanopipette. Cells were imaged with an IX83 Olympus microscope equipped with an X-Cite XYLIS illuminator (Excelitas Technologies, XT720S), ORCA-Fusion CMOS camera (Hamamatsu Photonics, C14440-20UP), and ×100 objective (Olympus, UPLXAPO100XO, NA 1.45).

### Quantitative PCR (qPCR) assay for mtDNA detection from individual mitochondria isolated by SICM

The template plasmid was constructed as follows; a part of mitochondrial DNA encoding 12S rRNA (chrM: 484-1,018) was amplified from DNA extracted from NIH-3T3 cells (RIKEN BRC, RCB2767) and cloned into pBluescript II SK(-), followed by confirmation by DNA sequencing. The copy number of plasmids per microliter was calculated from the concentration measured by NanoDrop One (Thermo Scientific^TM^) and molecular weight of the plasmid. The template plasmid was diluted to 5×10^7^ copies/µL with nuclease-free water plus yeast RNA (Roche, 10109223001), and this solution was further diluted to 2.5, 25, 200, 1600, and 12800 copies/µL and used for generating a standard curve. Low-binding tips (BMBio, W200-RS, FastGene, FGF-20LA) and low-binding tubes (Eppendorf, 0030108434) were used for dilution.

All qPCR reactions were run on a LightCycler 96 (Roche, 05815916001) using a QuantiNova Probe PCR Kit (Qiagen, 208254) following the kit protocol. qPCR amplification experiments were carried out in a 20 µL reaction volume consisting of 2× QuantiNova Probe PCR Master Mix, 0.4 µM dual-color probe, 0.2 µM forward and reverse primers, and nuclease-free water. The qPCR primer pairs (Fw: 5’-CTACCTCACCATCTCTTGCTAAT-3’, Rv: 5’-TTGGCTACACCTTGACCTAAC-3’) and the probe (5’-HEX-ATACCGCCA-ZEN-TCTTCAGCAAACCCT-IABkFQ-3’) were purchased from Integrated DNA Technologies. After an initial denaturation cycle of 95°C for 2 min, 50 PCR cycles were performed (denaturation at 95°C for 5 s, and annealing/extension at 60°C for 5 s). The quantification cycle (Cq) values (the number of PCR cycles at which the fluorescence amplification curve of a sample intersects the threshold line) were calculated using the fit points method of the LightCycler 96 software. The Cq values of the samples were fitted to the standard curve to determine the copy number of mtDNA, and those with the copy number < 1 were defined as mtDNA negative.

### Correlative light and electron microscopy

Primary cultured neurons were grown on no. 1S gridded coverslip-bottom dishes (custom made, based on IWAKI 3922-035; the grid-patterned side of the coverslips faced the inside of the dishes), precoated with carbon by a vacuum coater (EM ACE600, Leica), plasma-treated with a plasma ion bombarder (PIB-10, Vacuum Device Inc.) and then coated with poly-D-lysine (1 mg/ml, Sigma). Cells were incubated in Neurobasal media (Invitrogen), containing B27 (1x), Glutamax (1x), and Hoechst 33258 (1:500, FUJIFILM Wako, 080-09981) 3 hours before imaging. Fluorescence imaging was conducted at 10 DIV using a Nikon Ti2 Eclipse microscope with an A1R confocal, a CFI Plan Apochromat Lambda D 100X Oil (NA 1.45), a laser unit (Nikon; LU-N4, 405, 488, 561, and 640 nm), and filters (450/50 nm, 525/50 nm, 595/50 nm, 700/75 nm for 405 nm, 488 nm, 561 nm, 640 nm laser, respectively). All equipment was controlled via NIS-elements software. Optical sectioning was performed at Nyquist for the longest wavelength. The resulting images were deconvoluted with NIS-elements (Nikon) and processed with NIS-elements (Nikon) or ImageJ (NIH). Immediately after the fluorescent imaging, the cells were fixed with 2% paraformaldehyde (Electron Microscopy Sciences) and 2.5% glutaraldehyde (Electron Microscopy Sciences) in 0.1 M phosphate buffer (PB, pH 7.4) at 37°C for 1 hour and then washed with PB. The cells were then fixed with 2.5% glutaraldehyde in PB for 1 day at 4°C. After washing with PB, the cells were post-fixed with 1% OsO_4_ (Electron Microscopy Sciences), 1.5% potassium ferricyanide (FUJIFILM Wako) in PB for 30 minutes. After being rinsed 3 times with H_2_O, the cells were stained with 1% thiocarbohydrazide (Sigma-Aldrich) for 5 minutes. After being rinsed with H_2_O three times, the cells were stained with 1% OsO_4_ in H_2_O for 30 minutes. After being rinsed with H_2_O two times at room temperature and three times with H_2_O at 50°C, the cells were treated with Walton’s lead aspartate (0.635% lead nitrate (Sigma-Aldrich), 0.4% aspartic acid (pH 5.2, Sigma-Aldrich)) at 50°C for 20 minutes. The cells dehydrated with an ascending series of ethanol (10 minutes each in 50%, 70%, 90%, 95% ethanol/H_2_O, 100% ethanol four times on ice) were embedded in epoxy resin (Plain Resin, Nisshin EM), prepared by mixing 5.85 g of Plain Resin A, 5.85 g of Plain Resin B, and 0.21 g of Plain Resin C, by covering the gridded glass with a resin-filled beam capsule. Polymerization was carried out at 70°C for 95 hours. After polymerization, the gridded coverslip was removed and the resin block was trimmed to a square of about 1 mm. The block was sectioned using an ultramicrotome (EM UC7, Leica) equipped with a diamond knife (Ultra JUMBO 45 degree, DiATOME) to cut 50 nm thick sections. The serial ultra-thin sections were collected on the cleaned silicon wafer strip and imaged with a scanning electron microscope (JSM-IT800SHL; JEOL). The serial sections were imaged by a field emission scanning electron microscope (JSM-IT800SHL; JEOL) with the Array Tomography Supporter software (System in Frontier). Imaging was done at 1 kV accelerating voltage, 5 kV specimen voltage, 5,120 × 3,840 frame size, 6-mm working distance, 51.2 × 38.4-μm field of view and 5.33-μs dwell time, using a Scintillator Backscattered Electron Detector in Beam Deceleration mode. The final pixel size was a 10 nm square. The images taken by confocal microscopy were processed with ImageJ (NIH). The electron micrographs were aligned using the Linear Stack Alignment with Scale Invariant Feature Transform (SIFT) Plugin, implemented in ImageJ (NIH). Mitochondria, cell area, and cristae in electron micrographs were manually annotated. Reconstructing segmented images of the electron micrographs to 3-dementional images and overlaying it with fluorescence images were conducted using Imaris software (Bitplane).

### Investigation of electron microscopy images

Publicly accessible (42) Focused Ion Beam-Scanning Electron Microscopy (FIB-SEM) images of mouse somatosensory cortex layer 1 were analyzed. First, the deep learning-based mitochondrial segmentation tool, Empanada (72) was applied to the FIB-SEM dataset to infer all mitochondrial areas. After 3D reconstruction of the inferred mitochondrial areas, mitochondria fully contained within the dataset and localized to presynaptic boutons of excitatory neurons were selected and manually proofread to achieve precise extraction of mitochondrial regions. The extracted mitochondria were then visualized in three dimensions using Imaris version 9.6.0 (Bitplane).

### Quantification and Statistical Analysis

All statistical analysis and graphs were performed/created in Graphpad’s Prism 9. Statistical tests, p values, and (n) numbers are presented in the figure legends. Normality of data distribution was tested using D’Agostino & Pearson’s omnibus normality test. We applied non-parametric tests when data from groups tested deviated significantly from normality. All analyses were performed on raw imaging data without any adjustments. Images in figures have been adjusted for brightness and contrast (identical for control and experimental conditions in groups compared).

### Resources

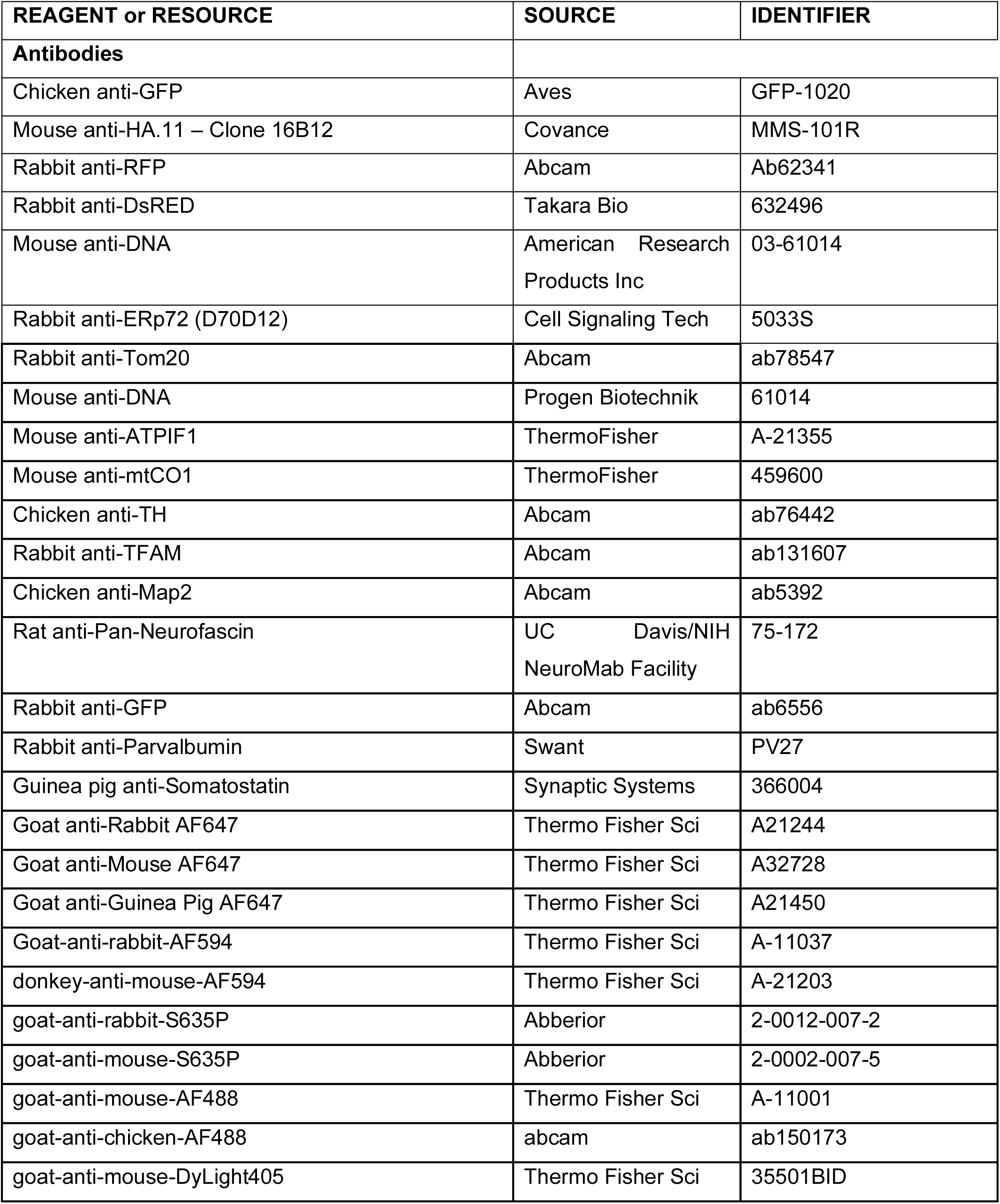
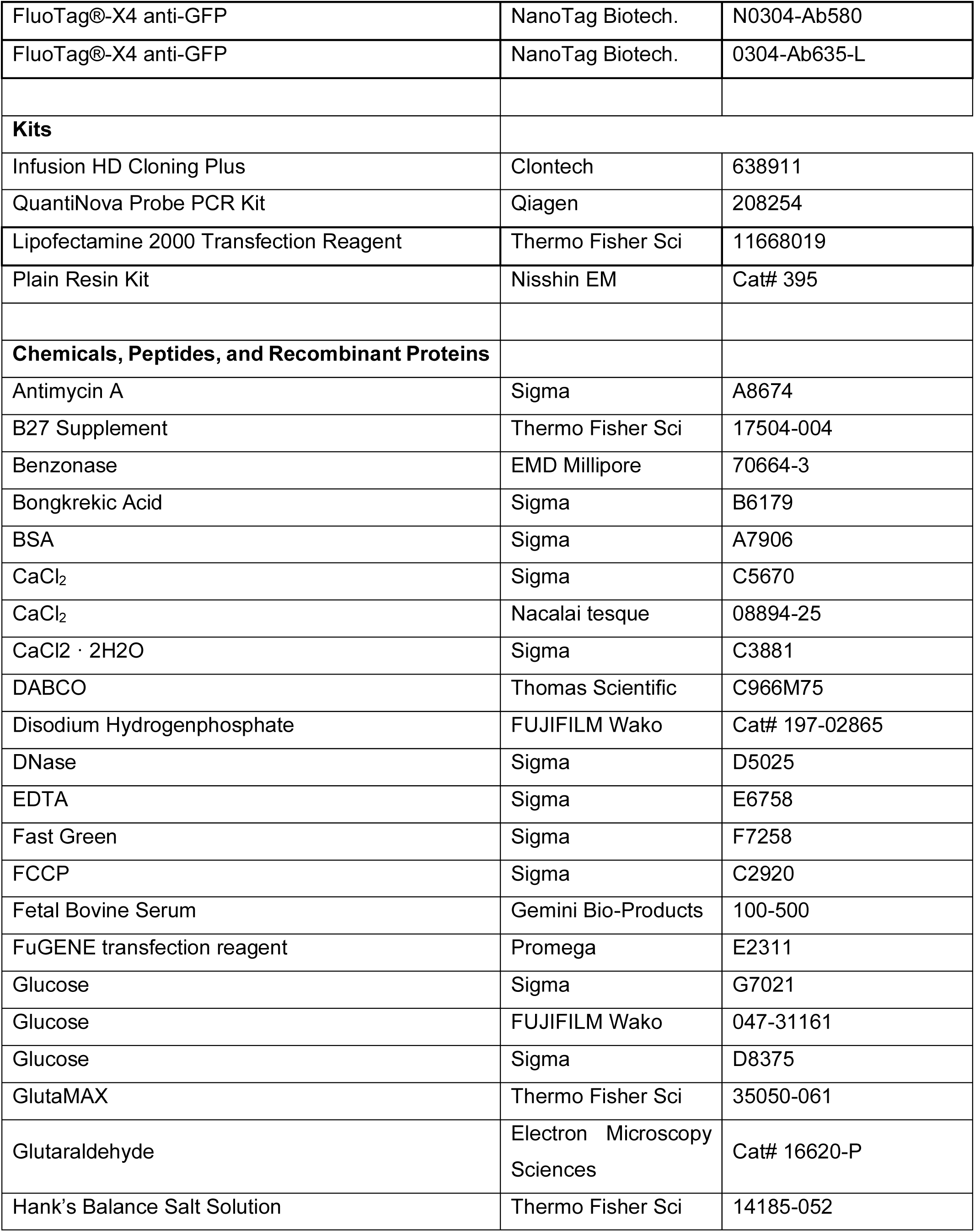
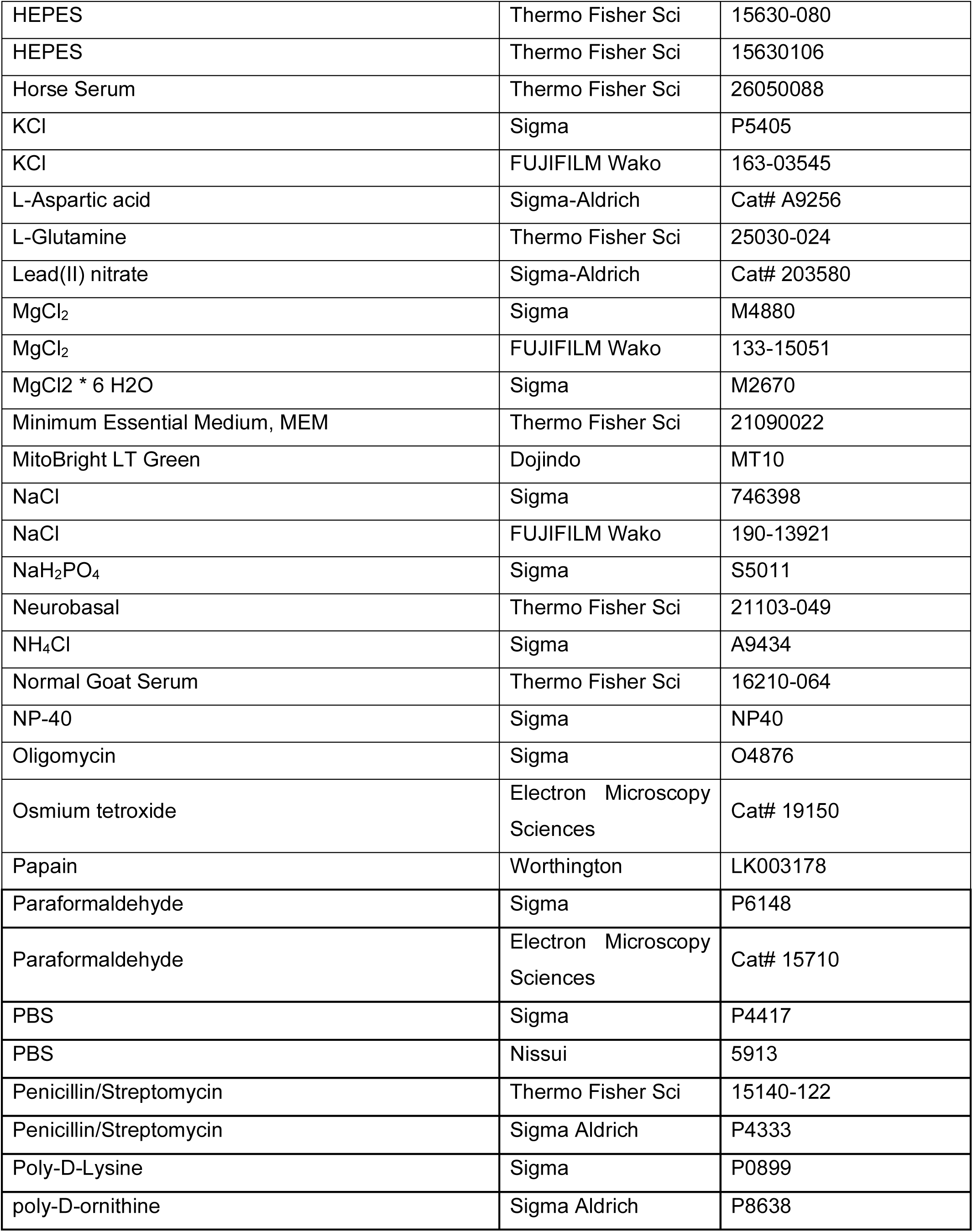
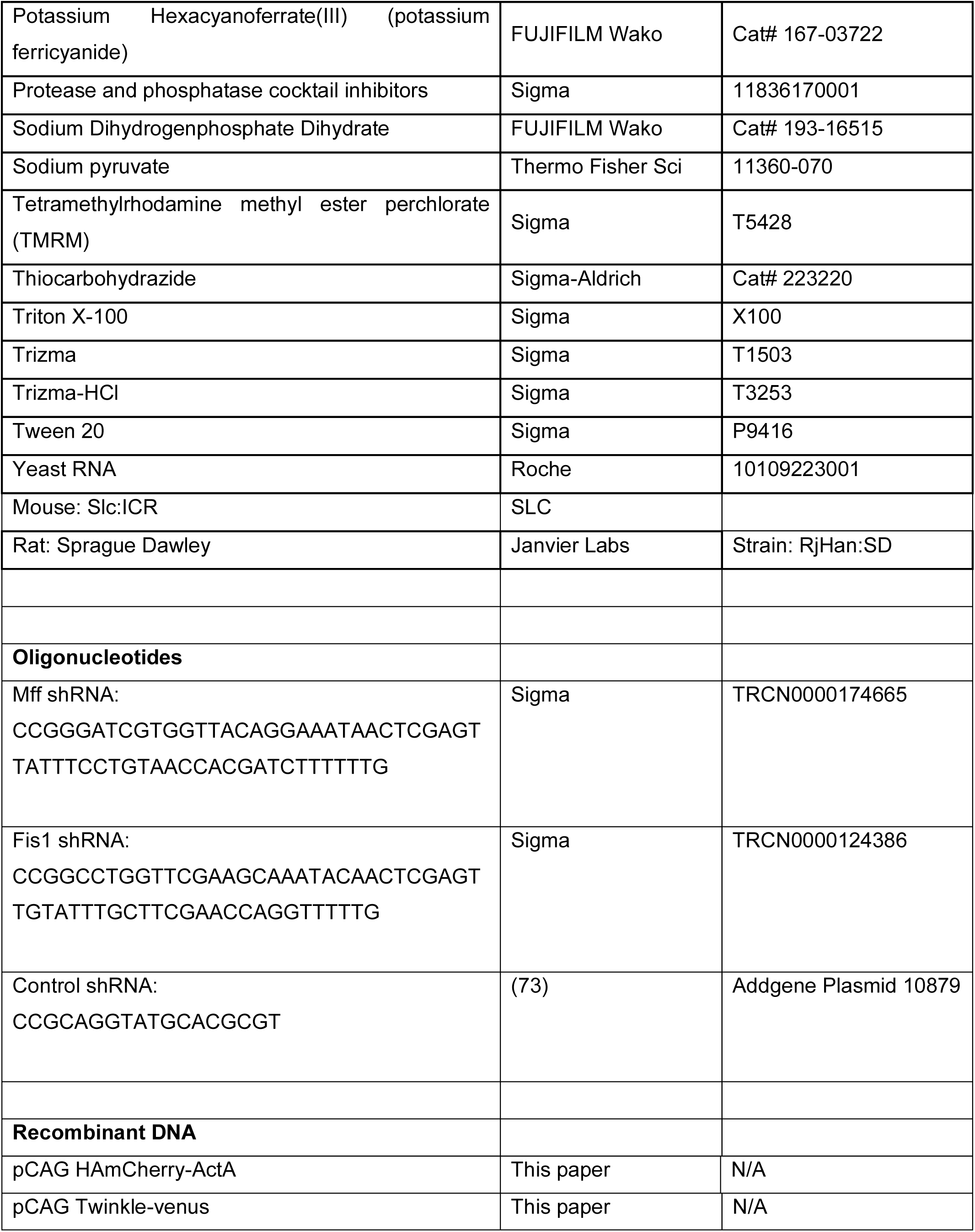
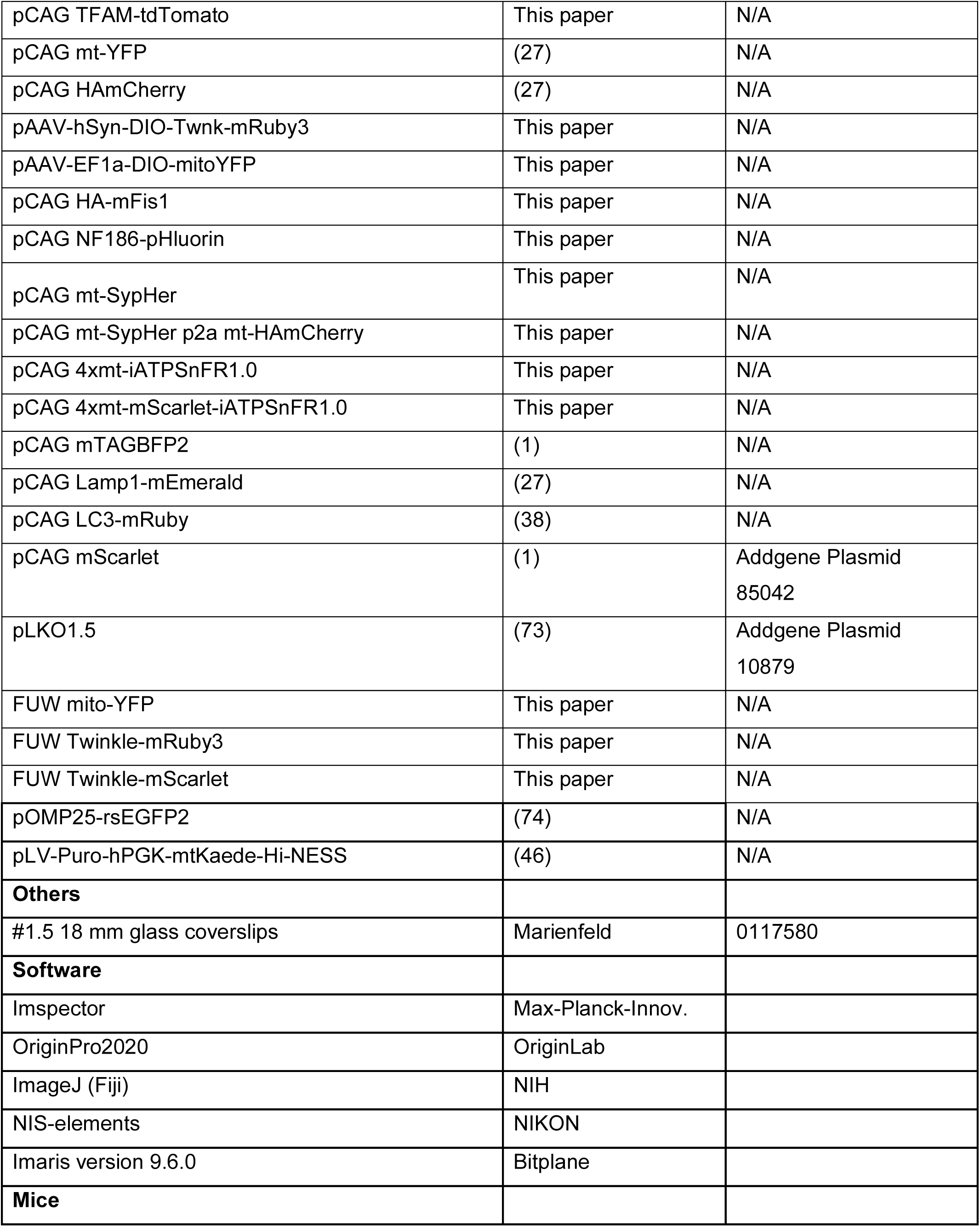
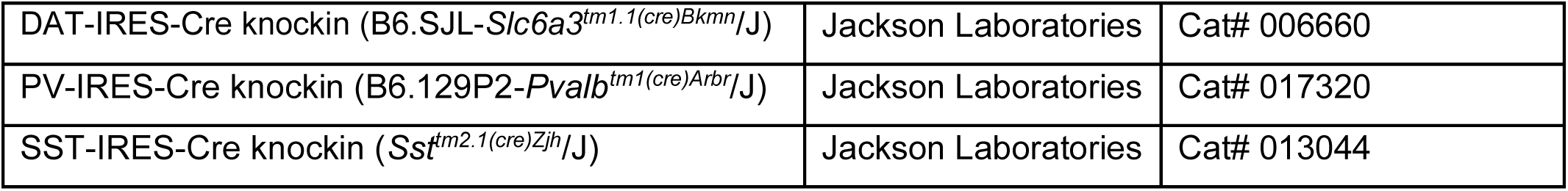

### Primary rat hippocampal neuron cultures

For the results shown in Figure 2, experiments were performed in Ilaria Testa laboratory in accordance with animal welfare guidelines set forth by Karolinska Institute and were approved by Stockholm Swedish Board of agriculture for Animal Research. Rats were housed with food and water available ad libitum in a 12h light/dark environment.

Primary hippocampal cultures were prepared from embryonic day 18 (E18) Sprague Dawley rat embryos. The pregnant mothers were sacrificed with CO_2_ inhalation and aorta cut; brains were extracted from the embryos. Hippocampi were dissected and mechanically dissociated in Minimum Essential Medium, MEM, (Thermo Fisher Scientific). 40 x 10^3^ cells per well were seeded in 12 well plates on a poly-D-ornithine (Sigma Aldrich) coated #1.5 18 mm glass coverslips (Marienfeld, ) and let them attach in MEM with 10% Horse Serum (Thermo Fisher Scientific), 2 mM L-Glut (Thermo Fisher Scientific) and 1mM Sodium pyruvate (Thermo Fisher Scientific), at 37°C at an approximate humidity of 95 – 98% with 5% CO2. After 3 hours the media was changed to Neurobasal Medium (Thermo Fisher Scientific) supplemented with 2% B-27 (Thermo Fisher Scientific), 2 mM l-Glutamine and 1% Penicillin-Streptomycin (Sigma Aldrich). The cultures were kept at 37°C at an approximate humidity of 95 – 98% with 5% CO2 for up to 24 days. Medium was changed twice per week. The experiments were performed on cultures starting from DIV7 up to DIV16.

### Immunostaining of neuronal culture for STED imaging

For data presented in Figure 2, rat neuronal cultures were washed in Artificial Cerebrospinal Fluid (ACSF) solution and then fixed in pre-warmed 4% paraformaldehyde (PFA) in phosphate buffered saline (PBS; pH 7.4) at RT for 15 minutes and permeabilized for 5 minutes in 0.1% Triton-X-100 buffer and blocked with 5% bovine serum albumin (BSA) (Sigma-Aldrich) in PBS, for 30 minutes at RT. Incubation with primary and secondary antibodies was performed in PBS solution for 1 hour at RT. Samples were mounted in custom-made Mowiol mounting media, supplemented with DABCO (Thomas Scientific).

Primary antibodies are listed and used as follows: anti-Tom20 (abcam, 1:200); anti-DNA (PROGEN Biotechnik, 1:100); anti-TFAM (abcam, 1:200); anti-Map2 (abcam, 1:2000); anti-Pan-Neurofascin (UC Davis/NIH NeuroMab Facility, 1:100), anti-GFP (abcam, 1:200). Secondary antibodies are listed and used as follows: Goat-anti-rabbit-AF594 (Thermo Fisher Scientific, 1:200 dilution), donkey-anti-mouse-AF594 (Thermo Fisher Scientific, 1:200 dilution); goat-anti-rabbit-S635P (Abberior, 1:200 dilution), goat-anti-mouse-S635P (Abberior,1:200 dilution); goat-anti-mouse-AF488 (Thermo Fisher Scientific, 1:200 dilution); goat-anti-chicken-AF488 (abcam, 1:200); goat-anti-mouse-DyLight405 (Thermo Fisher Scientific; 1:100), FluoTag®-X4 anti-GFP, (NanoTag, N0304-Ab580 or N0304-Ab635P-L).

### Neuron transfection and live cell staining for STED imaging

To stain the mitochondria outer membrane, neurons were transfected with OMP25-rsEGFP2 plasmid (74) using Lipofectamine 2000 Transfection Reagent (Thermo Fisher Scientific), according to the instructions of the manufacturer. To stain the AIS neuronal cultures were incubated for 5 minutes at RT with anti-pan-neurofascin primary antibody (UC Davis/NIH NeuroMab Facility, 1:100), subsequently washed three imes with ACSF buffer, then incubated with goat-anti-mouse-DyLight405 secondary antibody (Thermo Fisher Scientific; 1:100) for 5 minutes at 37°C and finally washed three times in ACSF buffer.

### STED imaging

STED images were recorder with a custom-built STED setup, previously described (30). Excitation of the dyes was done with pulsed diode lasers; at 561 nm (PDL561, Abberior Instruments), 640 nm (LDH-D-C-640, PicoQuant) and 510 nm (LDF-D-C-510, PicoQuant). A laser at 775 nm (KATANA 08 HP, OneFive) was used as the depletion beam, which was split into two orthogonally polarized beams that were separately shaped to a donut and a top-hat respectively in the focal plane using a spatial light modulator (LCOS-SLM X10468-02, Hamamatsu Photonics). The laser beams were focused onto the sample using a HC PL APO 100×/1.40 Oil STED White objective (15506378, Leica Microsystems), through which the fluorescence signal was also collected. The images were recorded with a 561nm excitation laser power of 8–20 µW, a 640 nm excitation laser power of 4–10 µW and a 775 nm depletion laser power of 128 mW, measured at the first conjugate back focal plane of the objective. Two-color STED imaging was done in a line-by-line scanning modality. The pixel size was set between 20 and 30 nm with a pixel dwell time of 50 µs.

Volumetric data was acquired by successive 2D-STED imaging of nucleoids was recorded with a voxel size for xyz volumes was set to 25 × 25 × 200 nm^3^ (x-y-z). The pixel dwell time was set at either 30 or 50 µs. The spatial resolution of the 2D-STED modality microscope was measured on a calibration sample where sparse antibodies coupled to the STAR 635P dye were attached to the cover glass. The line profiles were extracted and fitted with a Lorentzian function (75), from which the full width at half maximum (FWHM) was extracted. The distribution of the FWHM values was peaking at 42 ± 9 nm (mean ± SD, n = 50) (**Extended Data Fig. 3**).

### Image analysis of super-resolution (STED) microscopy

The images were processed and visualized using the ImSpector software (Max-Planck Innovation) and ImageJ(76). When necessary, images and movies were deconvolved using the Richardson-Lucy algorithm, implemented in Imspector. The PSF was modelled as a Gaussian function and the FWHM was chosen to be 40 nm. The regularization parameter and number of iterations were varied depending on the quality of the output image. The regularization parameter was set as either 10^-5^ while the number of iterations was chosen up to a maximum of 5. Brightness and contrast were linearly adjusted for the entire images. The data were then analysed, fitted and visualized with the software OriginPro2020 (OriginLab). To calculate the frequency of mitochondria without and with (1-6+) nucleoids in axons and dendrites, a analysis pipeline was developed: (i) the binary maps of soma, dendrites and axons were generate in ImageJ, based on the Map2 and Neurofascin staining; (ii) the mitochondria binary map was generated based on mitochondria staining; (iii) To measure the distribution of nucleoids per mitochondria along the axonal length, a semi-automatic pipeline was set-up. It consists of the following steps: (1) Binarize image of mitochondria and obtain center points of each mitochondria; (2) Detect maxima in TFAM image (glass-to-glass adjusted thresholding) and extract the positions; (3) Count the number of maxima per binarized mitochondria area; (4) Manually segment axon from AIS image and binarize; (5) Make a geodesic distance transform from manually selected seed point closest to the soma; (6) Check distance from each mitochondria center to the soma in the geodesic distance map. For comparison of distributions of parameters, the Kolmogorov-Smirnov test was chosen (KS-test). This choice was made due to the non-normal nature of the observed distributions.

### Fluorescent In Situ Hybridization (FISH) detection of mitochondrial DNA (mtDNA) and mtDNA encoded mRNA

For results shown in Figure 4, the detection of mitochondrial DNA and mtDNA transcript mRNA was conducted by single molecule fluorescent in situ hybridization (FISH) of specific targeting probes with a modified version of the commercial RNAscope Multiplex Fluorescent v2 Assay Protocol. The RNAscope targeting scheme utilizes a pair of gene-specific double Z probes (ZZ) that together bind a contiguous region of approximately 50 bases on the target sequence. Each Z probe includes a spacer region and a 14-base tail sequence that, when aligned next to the other probe’s tail sequence, generates a 28 -base binding site for the preamplifier oligonucleotide (77). The preamplifier contains 20 binding sites for the amplifier, which, in turn, contains 20 binding sites for the label probe, allowing signal amplification similarly to the previously described branched DNA scheme (78). The double Z probe strategy confers high target specificity because amplification is dependent on both Z probes localizing to their target sequences to generate the 28-base landing site. For the two protocols employed here to detect mitochondrial DNA and mRNA transcripts, samples were pretreated with RNAse or DNAse, respectively, to exclusively target the molecule of interest.

#### Detection of mtDNA using DNAscope

The modified protocol used for labeling DNA of mitochondrial genes in cultured mouse cortical neurons is referred to here as “DNAscope” as the resulting signal corresponds to the mitochondrial DNA instead of mitochondrial RNA transcripts. For Figure 4a-e, ex-utero cortical electroporation was performed on E15.5 mouse embryos in order to target the neural progenitors generating layer 2/3 PNs, immediately dissociated and plated at 100,000 cell density on 35 mm MatTek Dishes, cultured for either 14 or 21 days, and fixed in 4% paraformaldehyde (PFA; EMS; 15710-S) for 15 minutes, followed by three 5 minute washes with PBS (1X PBS; Gibco™ 10010049). The cells were then treated with RNAscope® Hydrogen Peroxide Reagent for 10 minutes at 23°C to 25°C and washed twice with UltraPure distilled water (Invitrogen; 10977015), followed by digestion in a 1:15 dilution of RNAscope® Protease III Reagent in PBS at 23°C to 25°C and two washes in PBS. Inclusion of a subsequent incubation step with RNAse cocktail to clear mRNA signal was the first modification to the RNAscope protocol. Immediately following hydrogen peroxide and protease treatment, the cells were incubated with the RNAse cocktail (RNase A at 20 U/mL; RNase T1 at 800 U/mL; Fisher Scientific) in PBS for 30 minutes at 37°C inside a HybEZ hybridization oven (ACD; PN 321710). These steps constituted the pretreatment steps for fixed cell culture samples. Ethanol dehydration was not used as this step would damage the critical fluorescent protein that labeled mitochondria in electroporated neurons. The samples were then washed twice with PBS and then incubated with pre-warmed target probes (20 nmol/L of each oligo probe) overnight at 40°C inside the HybEZ hybridization oven. The mitochondrial Cytochrome B sequence (CY-B) was targeted with RNAscope® Probe- Mm-mt-Cytb (ACD;Cat No. 517301), and the mitochondrial Cytochrome C oxidase subunit I sequence (CO1) was targeted with RNAscope® Probe- Mm-mt-Co1-C2 (ACD;Cat No. 517121-C2). The second modification to the RNAscope protocol involved extending the primary target probe incubation step to overnight (18-21 hours) at 40°C instead of 2 hours at 40°C inside the HybEZ hybridization oven. These two modifications have been previously implemented for targeting viral DNA (79). The option of a 60°C DNA denaturation step previously used for targeting viral DNA (79) was excluded as it would also denature the critical fluorescent protein that labeled mitochondria in electroporated neurons. After the overnight target probe hybridization, the samples were incubated at 40°C in Amplifier 1 (preamplifier) (2 nmol/L) in hybridization buffer B (20% formamide, 5× SSC, 0.3% lithium dodecyl sulfate, 10% dextran sulfate, blocking reagents) for 30 minutes; Amplifier 2 (2 nmol/L) in hybridization buffer B at 40°C for 15 minutes; and Amplifier 3 (label probe) (2 nmol/L) in hybridization buffer C (5× SSC, 0.3% lithium dodecyl sulfate, blocking reagents) for 15 minutes. After each hybridization step, slides were washed with wash buffer (0.1× SSC, 0.03% lithium dodecyl sulfate) two times at room temperature (80). Chromogenic detection was performed utilizing a horseradish peroxidase (HPR) construct specific to each gene-dedicated imaging channel and a fluorescent Opal reagent of choice. CY-B was stained with Opal 520 Reagent (Perkin Elmer, FP1487001KT), and CO1 was stained with Opal 570 Reagent (Perkin Elmer, FP1488001KT). Each Opal reagent dye was diluted 1:1500 in RNAscope® Multiplex TSA Buffer. Coverslips were mounted onto slides in Fluoro-Gel (EMS; 17985-10) and analyzed at 60x magnification using a Nikon A1 confocal microscope.

#### Detection of mtDNA-encoded mRNA using mt-RNAscope

Labeling mRNA transcripts of mtDNA-encoded CytB and Atp6 was performed using a further modified version of the RNAscope® Multiplex Fluorescent v2 Assay protocol. We performed EUE on E15 mouse embryos to express mitoYFP (green matrix-targeted mitochondrial marker), dissociated as described above and maintained in culture for 18-21 DIV. Cultures were fixed using 4% PFA and 4% sucrose (Boston BioProducts; IBB-255) for 10 min, followed by 2 washes in PBS and cells were then permeabilized using 0.1% Triton X-100 (Sigma Aldrich; 9036-19-5) for 10 min and washed twice for 5 min in PBS. Following these steps, cells were treated with RNAscope® Hydrogen Peroxide Reagent for 10 min at RT and washed twice with UltraPure distilled water. Both the Ethanol dehydration and RNAscope® Protease III steps were excluded from the protocol to preserve the mitochondrial marker fluorescence and overall organelle morphology. Instead of using an RNAse cocktail, cells were then treated with DNAse I (NEB; M0303S) for 30 minutes at 37°C in order to exclusively detect mRNA transcripts. Following this, the protocol was carried out as described in the RNAscope® Multiplex Fluorescent v2 Assay protocol. To detect *CytB* and *Atp6* RNA transcripts, a 1:2 dilution of the RNAscope® Probe-Mm-mt-Cytb Cat No.517301 and RNAscope® Probe-Mm-mt-Atp6 Cat No.544401 and a 1:3000 dilution of Opal 690 Reagent (Akoya; FP1497001KT) in RNAscope® Multiplex TSA Buffer were used. Following the RNAscope protocol, dishes were mounted and imaged in a W1-Yokogawa inverted spinning disk confocal microscope. The number of RNA foci overlapping with mitoYFP signal was quantified for dendritic and axonal mitochondria in image stacks using Fiji.

## Acknowledgments

We thank Pierre Vanderhaeghen, and members of the Lewis, Hirabayashi and Polleux labs for their comments on the manuscript. We thank David Ng (ZMBBI Molecular Tools Core Facility at Columbia University), Qiaolian Zhang, Joshua Weertman, and Kelsey Carter for excellent technical assistance.

## Funding

JSPS KAKENHI Grant Numbers JP19H03221 (YH), JP22H02716 (YH), JP22K18939 (YT), and JP22KJ1098 (YD)

AMED Grant numbers JP19dm0207082 (YH), JP21wm0525015 (YH, YT), JP 25wm0625318 (YH)

JST, PRESTO Grant Number JPMJPR16F7 (YH) and JPMJPR14FA (YT)

National Institute of Health- National Institute of Neurological Disorders and Stroke- R35 NS127232 (FP)

JST, FOREST Grand Number JPMJFR203K (YT)

JST, CREST Grant Number JPMJCR24T6 (YH, YT)

Human Frontier Science Program (HFSP) (RGP0028/2022) (YT)

National Institute of Health- National Institute of General Medical Sciences – R35 GM137921 (TLL)

ERC-CoG Inspire (IT, GC, JA)

The Leona M. & Harry B. Helmsley Charitable Trust (1903-03788) (JG)

NIH NCI 1DP2CA281605-01 (JG)

## Author contributions

Conceptualization: FP, TLL, YH

Methodology: YH, TLL, EZ, YD, DMV, JUJ, AMD, GC, JA, MB, MT, PC, TES, YT, JTG, IT, FP

Investigation: YH, TLL, EZ, YD, DMV, JUJ, AMD, SH, GC, MK, MS, SS, JA, MP, PK, AM

Visualization: YH, TLL, EZ, YD, DMV, JUJ, AMD, GC, SS, PK, YT, JTG, IT, FP

Funding acquisition: YH, TLL, YT, JLG, IT, FP

Project administration: YH, TLL, FP

Supervision: TLL, YH, JTG, IT, FP

Writing – original draft: TLL, YH, FP

Writing – review & editing: YH, TLL, EZ, YD, DMV, AMD, GC, PK, PC, TES, YT, JTG, IT, FP

## Competing interests

Authors declare that they have no competing interests.

## Data and materials availability

All data are available in the main text or the supplementary materials.

## Notes

### Competing Interest Statement

The authors have declared no competing interest.

### Summary of Updates

New work has been added to demonstrate the mtDNA phenotype in additional neuron types in vivo (Figure 3). Additional experiments were performed to show mtRNA and mtDNA encoded proteins are lacking in axonal mitochondria (Figure 4). mt-HINESS was used to confirm that new mitochondria entering processes are mostly lacking mtDNA (Figure 6). Author list updated; Supplemental files updated.

