## Supplemental Figures for "The majority of axonal mitochondria in mammalian neurons lack mitochondrial DNA and do not produce ATP"

**Supplementary Information and movies**

Hirabayashi, Lewis et al. 2025

Figs. S1 to S11

Movies S1-4

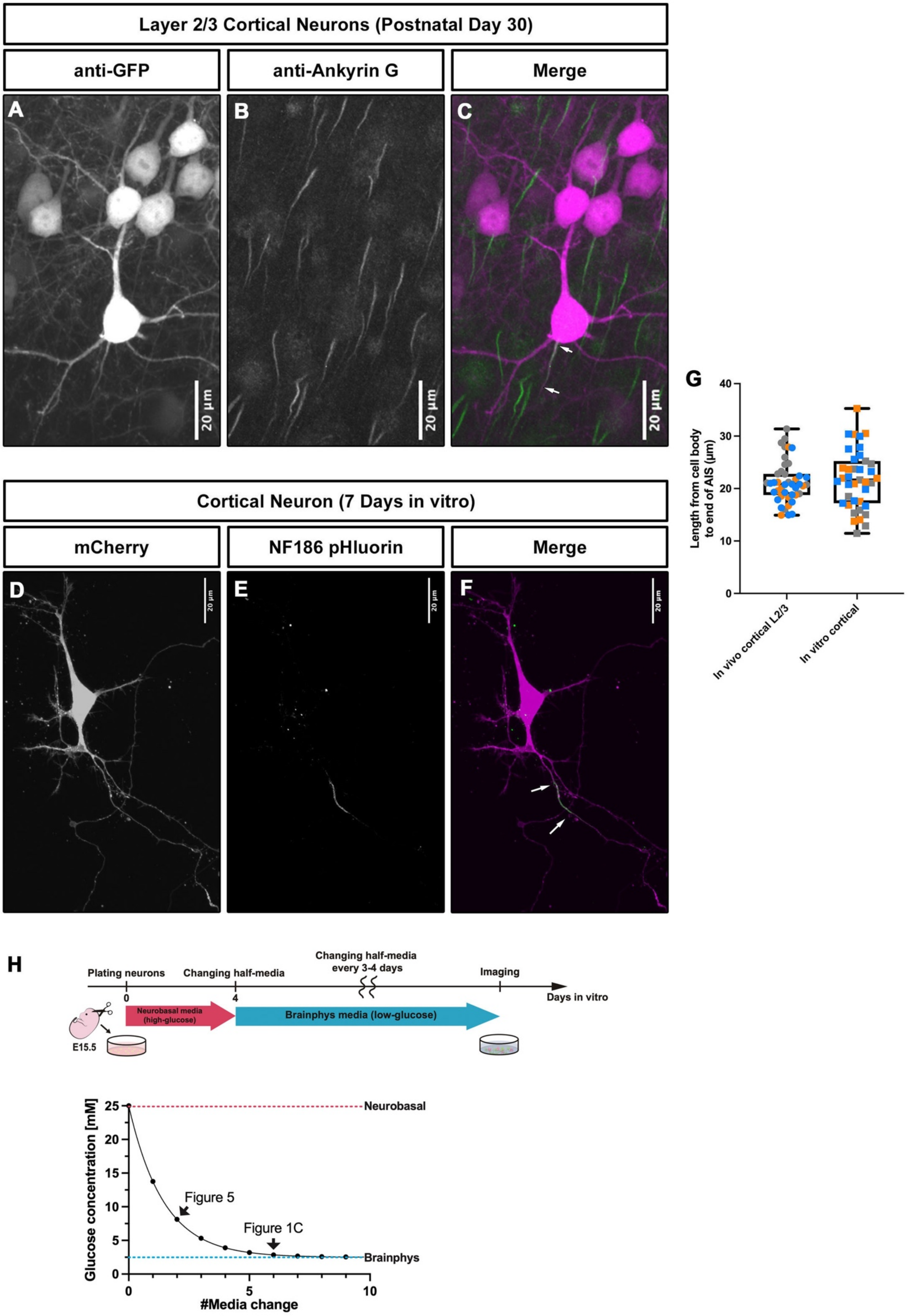

Layer 2/3 Cortical Neurons (Postnatal Day 30)
anti-GFP
anti-Ankyrin G
Merge
A
B
C
20 µm
Cortical Neuron (7 Days in vitro)
mCherry
NF186 pHluorin
Merge
D
E
F
20 µm
G
Length from cell body to end of AIS (µm)
0
10
20
30
40
In vivo cortical L2/3
In vitro cortical
H
Plating neurons
Changing half-media
Changing half-media every 3-4 days
Imaging
Days in vitro
0
4
Neurobasal media (high-glucose)
Brainphys media (low-glucose)
E15.5
Glucose concentration [mM]
#Media change
Neurobasal
Brainphys
Figure 5
Figure 1C
0
5
10
15
20
25

**Supplemental Figure 1. Measurement of axon initial segment (AIS) length in layer 2/3 CPNs *in vivo* and *in vitro*.** Related to Figure 1. (**A-C**) Representative maximum projection images of layer 2/3 cortical neurons labelled with GFP via *in utero* electroporation at E15.5, followed by perfusion and staining with anti-GFP and anti-Ankyrin G at P30. (**D-F**) Representative maximum projection images of a cultured cortical neuron labelled with cytoplasmic mCherry and Neurofascin (NF)186-phluorin via *ex utero* electroporation at E15.5, followed live confocal imaging at 7DIV. Arrows in panels c and f point to AIS. (**G**) Quantification of the length from the cell body to the end of the axon initial segment (AIS) for layer 2/3 cortical neurons *in vivo* and *in vitro* showing that in both cases the average length to the end of the AIS is ~ 20 microns. n = 43 neurons from 3 independent brains for a-c. n = 39 neurons from 3 independent cultures for d-f. Same coloured data points are from the same independent brain or culture. White arrows point to AIS. Scale bars 20μm. (**H**) (Top) Schema of the primary cortical neuronal culture under physiological glucose conditions: Mouse neurons were cultured in Neurobasal medium (25 mM glucose) for the first 4 DIV. At 4 DIV, half of the medium was replaced with BrainPhys medium (2.5 mM glucose), with subsequent half-medium changes every 3 to 4 days. (Bottom) Calculated glucose concentration throughout the culture period without accounting for glucose consumption by cultured cells. The mitochondrial isolation performed in Fig. 5 was performed after two medium exchanges when the glucose concentration dropped below 8 mM. The quantification of Twinkle+ mitochondria in Fig. 1c was performed after six medium exchanges when the glucose concentration had approached 2.5 mM.

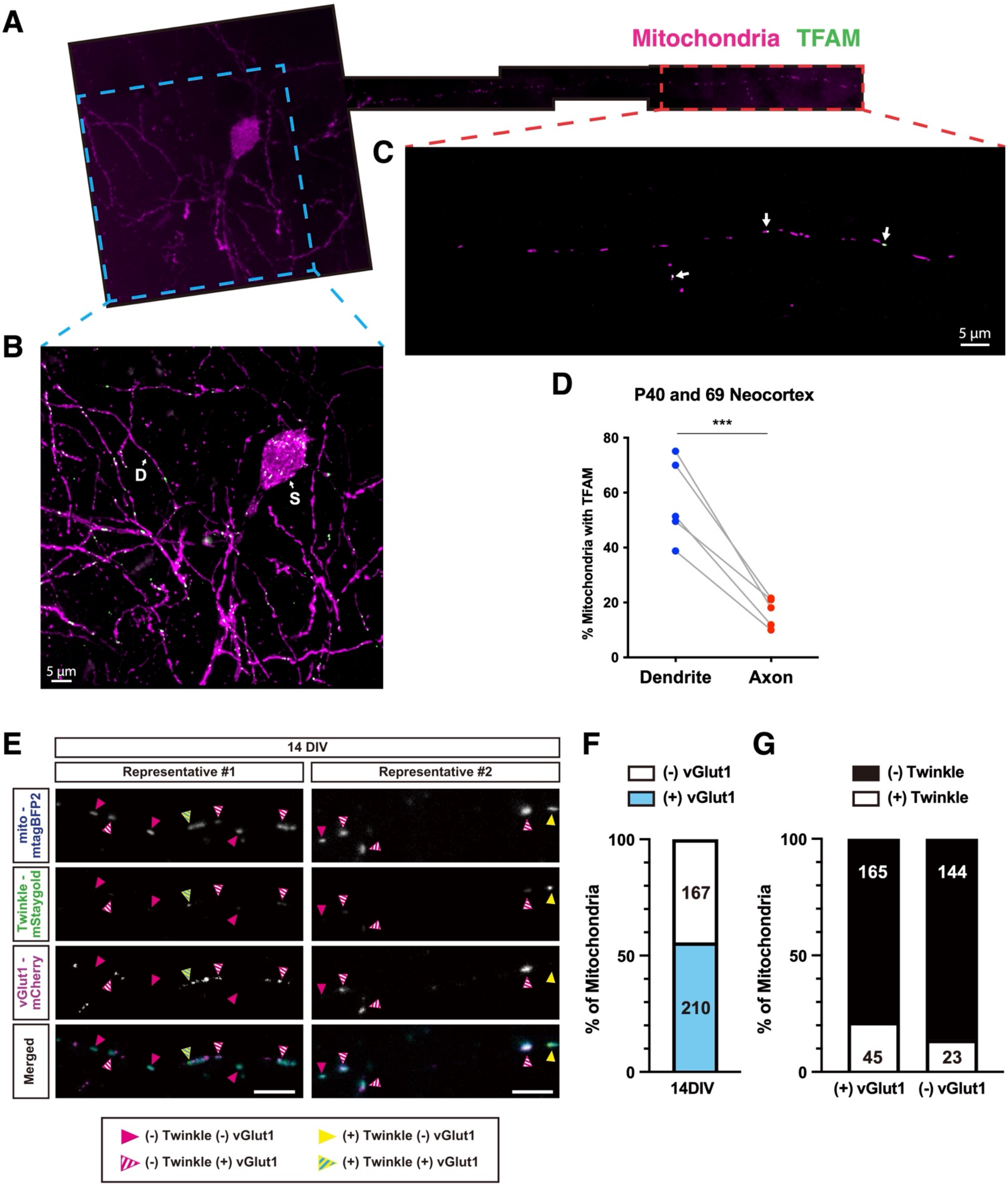
A
Mitochondria TFAM
B
D
S
5 µm
C
5 µm
D
P40 and 69 Neocortex
***
% Mitochondria with TFAM
80
60
40
20
0
Dendrite
Axon
E
14 DIV
Representative #1
Representative #2
mito - mtagBFP2
Twinkle - mStaygold
vGlut1 - mCherry
Merged
(-) Twinkle (-) vGlut1
(-) Twinkle (+) vGlut1
(+) Twinkle (-) vGlut1
(+) Twinkle (+) vGlut1
F
(-) vGlut1
(+) vGlut1
% of Mitochondria
100
50
0
167
210
14DIV
G
(-) Twinkle
(+) Twinkle
% of Mitochondria
100
50
0
165
144
45
23
(+) vGlut1
(-) vGlut1

**Supplemental Figure 2. A low fraction of axonal mitochondria contains mtDNA-associated protein TFAM *in vivo* and Twinkle *in vitro* independently of whether mitochondria are associated with presynaptic boutons or not.** Related to Figure 1. (**A**) Representative image of an optically isolated layer 2/3 CPN expressing TFAM-tdTomato (green) and mitochondrial matrix-targeted mt-YFP (magenta) *in vivo* at P40. (**B**) High magnification of mitochondria in the soma (S) and the dendrites (D). (**C**) High magnification of the proximal part of the axon with the few mitochondria containing TFAM labeled by arrows. Scale bar: 5µm. (**D**) Percentage of mitochondria positive for TFAM in the axon or dendrites of layer 2/3 CPNs in vivo. Numbers of mitochondria counted are shown in each column. 5 neurons from 2 mice at each of P40 and P69 were used for quantifications. *** p=0.0004 by paired t-test. (**E-G**) Primary cultured neurons expressing a mitochondrial marker (mito-mtagBFP2), an mtDNA marker (Twinkle-mStaygold), and a presynaptic marker (vGlut1-mCherry) were imaged at 14 DIV. (E) Representative images of axonal segments. Magenta or yellow arrowheads indicate mitochondria marked negative or positive for Twinkle, respectively. Striped arrowheads indicate mitochondria localized at presynaptic boutons. (F) Percentage of mitochondria localized at presynaptic boutons ((+) vGlut1). (G) Percentages of presynaptic ((+) vGlut1) or non-presynaptic ((-) vGlut1) mitochondria that are positive (white bars; (+) Twinkle) or negative (black bars: (-) Twinkle) for Twinkle. The numbers of mitochondria counted are indicated in each column. Scale bars: 5 µm.

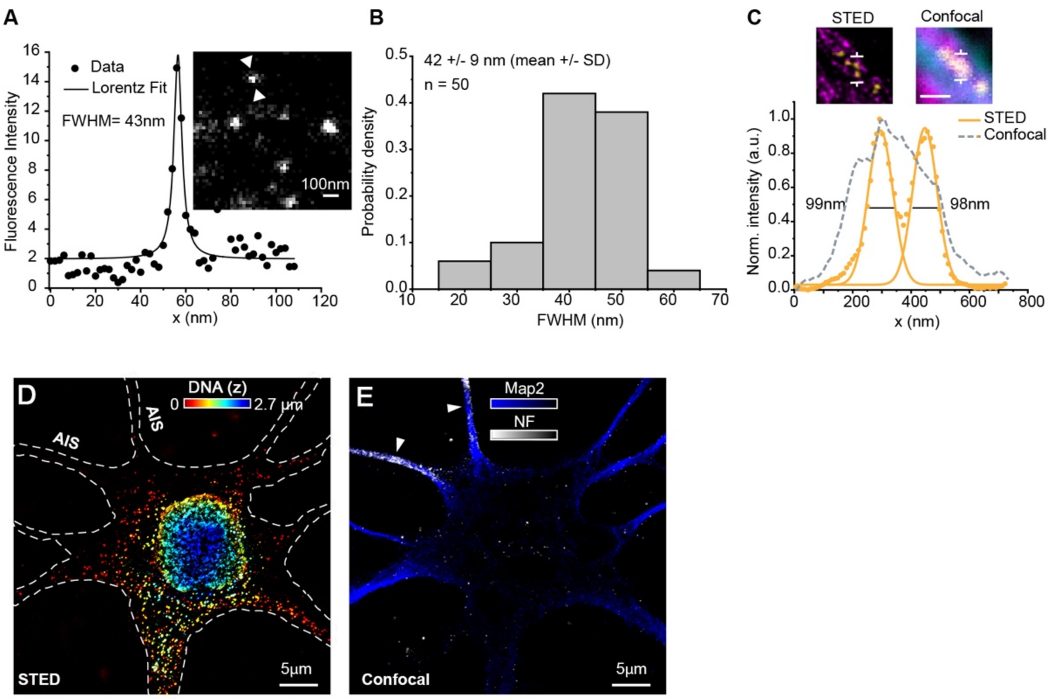


**Supplemental Figure 3. Super-resolution STED imaging of mitochondria nucleoids in primary rat hippocampal neurons.** Related to Figure 2.

(**A**) Measurement of STED resolution using STAR 635P-conjugated antibodies. Line profile over single antibody as indicated in the insert, and Lorenzian fit of this line profile. (**B**) Distribution of the FWHM of Lorenzian-fitted line profiles of antibodies, representing the resolution of the STED microscope: 42 ± 9 nm (mean ± SD), n = 50 from one experiment. (**C**) upper panels: inset of Fig. 1B where single nucleoids are distinguishable in STED, but not in the confocal comparison; bottom panel: line profiles measured at the indicated signs for STED (yellow dots and Gaussian fit) and for confocal (grey dotted line). Scale bar 500nm. (**D**) 2D-STED volumetric image showing the spatial distribution of nucleoids (immunostained for DNA) across the cell soma. Nucleoids are barely visible along the AIS, identified via the corresponding NF staining (arrowheads in E). (**E**) Confocal image showing the dendritic marker Map2 (blue) and the AIS marker neurofascin (NF) labelling the AIS (arrowheads).

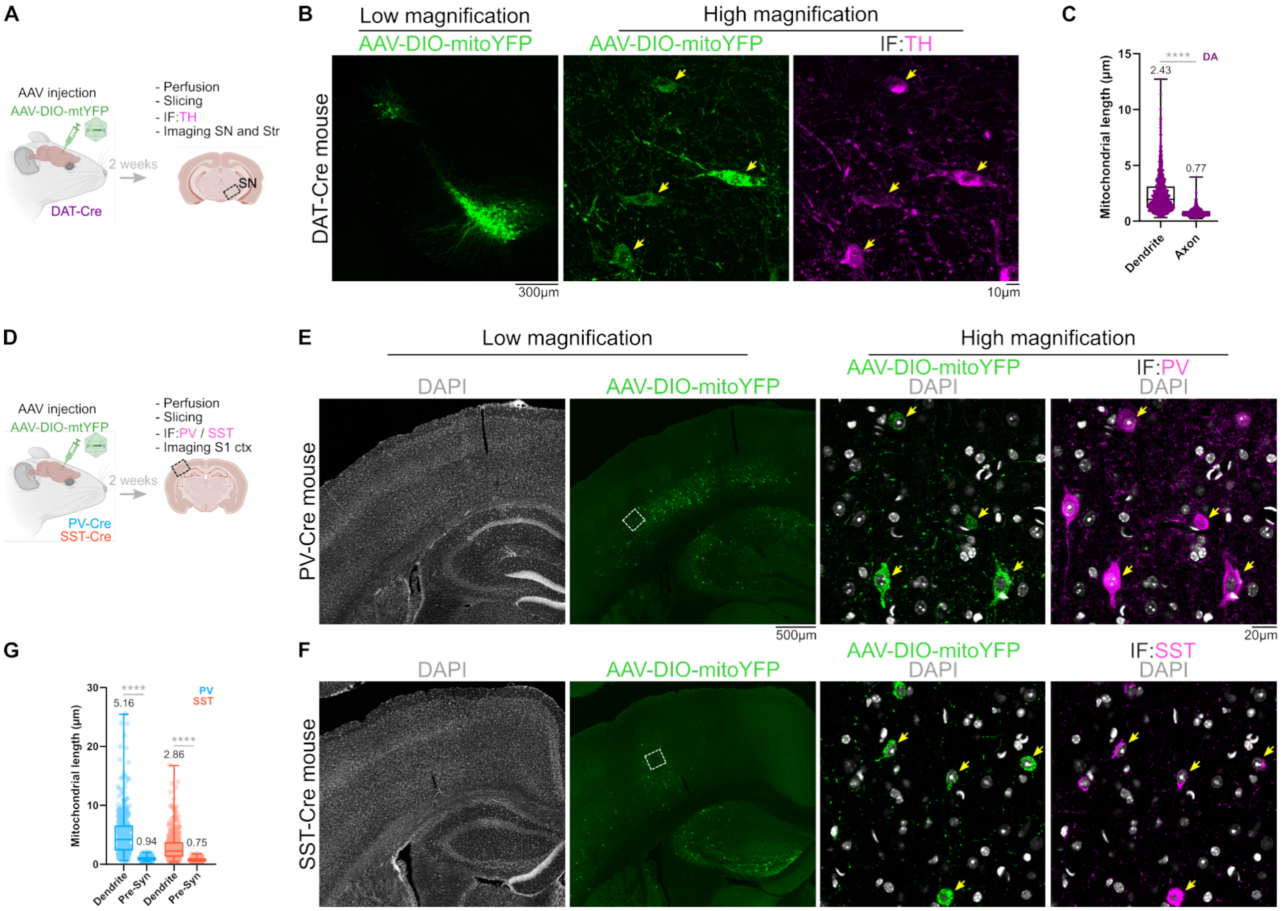


**Supplemental Figure 4. Validation of genetic approaches to target expression of Cre-dependent AAV in dopaminergic neurons of substantia nigra pars compacta (SNc), cortical PV+ interneurons (PV-INs) and SST+ interneurons (SST-INs) of adult mice.** Related to Figure 3.

(**A**) Schematic representation of the experimental pipeline used in Figure 3b-d, except brain slices were stained for TH to determine the specificity of AAV expression in SNc of DAT-Cre mice. (**B**) Representative low- and high-magnification images showing the expression of mitoYFP (green) in TH-positive neurons (magenta). Arrows indicate double-positive neurons. (**C**) Quantification of mitochondrial length in the dendritic and axonal compartments of DA neurons. n axonal mitochondria = 1016, n dendritic mitochondria = 1510 from 6 mice. p<0.0001 (****) by Mann-Whitney for non-parametric data. (**D**) Schematic representation of the experimental pipeline employed in Figure 3f-h, except brain slices were stained for PV or SST to determine the specificity of AAV expression in the PV-Cre and SST-Cre animals, respectively. (**E-F**) Representative low- and high-magnification images showing the expression of mitoYFP (green) in PV (**E**) or SST (**F**) positive INs (magenta). Arrows indicate double-positive neurons. (**G**) Quantification of mitochondrial length in the dendritic and pre-synaptic compartments of PV- and SST-INs from experiments in Figure 3g-h and 3m-n. Each dot represents individual mitochondria, with 320-810 mitochondria from 8-10 mice analyzed per group. p<0.0001 (****) by Mann-Whitney for non-parametric data.

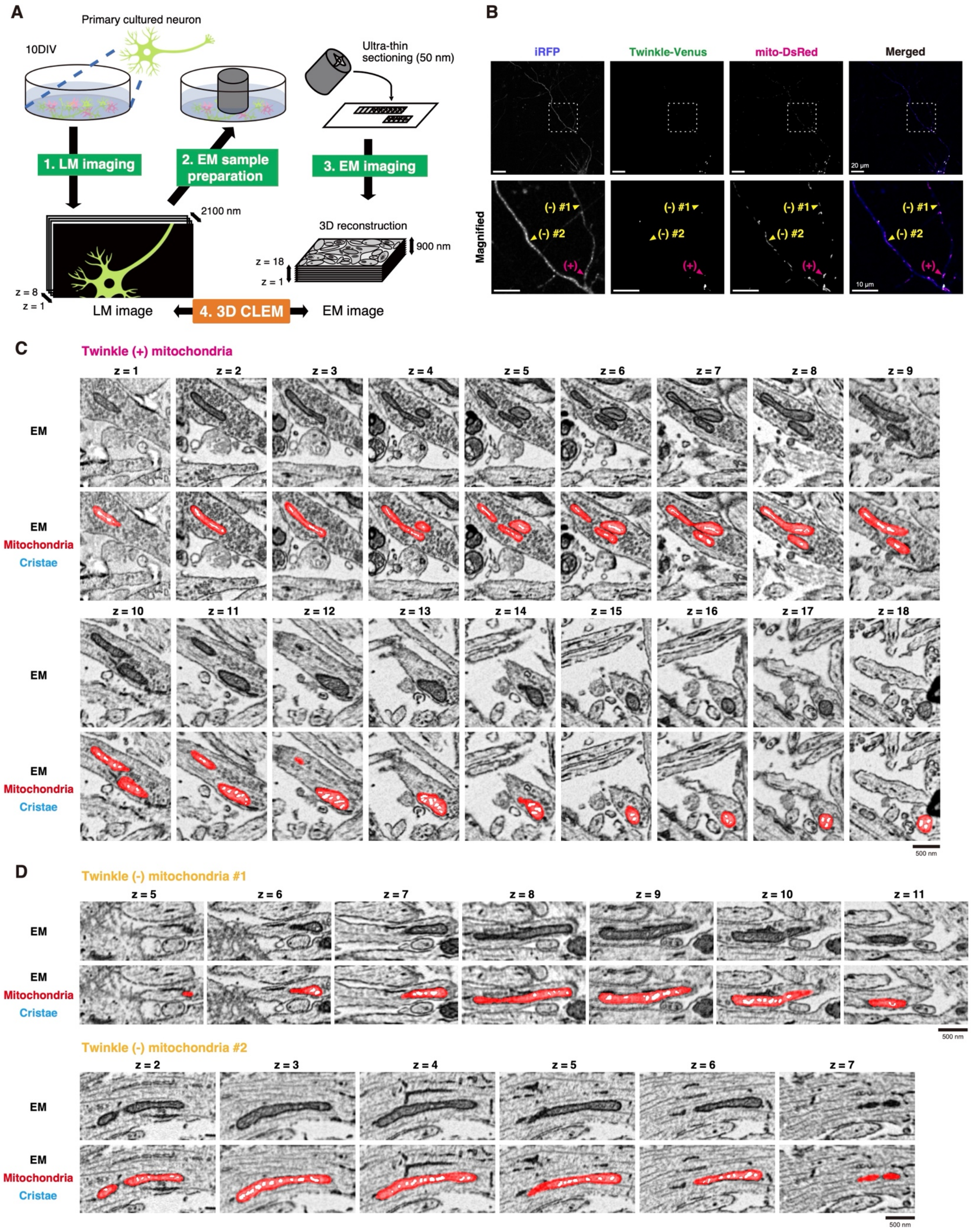

A
Primary cultured neuron
10DIV
Ultra-thin sectioning (50 nm)
1. LM imaging
2. EM sample preparation
3. EM imaging
2100 nm
3D reconstruction
900 nm
z = 18
z = 1
z = 8
z = 1
LM image
4. 3D CLEM
EM image
B
iRFP
Twinkle-Venus
mito-DsRed
Merged
20 µm
Magnified
(-) #1
(-) #2
(+)
10 µm
C
Twinkle (+) mitochondria
EM
EM
Mitochondria
Cristae
z = 1
z = 2
z = 3
z = 4
z = 5
z = 6
z = 7
z = 8
z = 9
z = 10
z = 11
z = 12
z = 13
z = 14
z = 15
z = 16
z = 17
z = 18
500 nm
D
Twinkle (-) mitochondria #1
z = 5
z = 6
z = 7
z = 8
z = 9
z = 10
z = 11
500 nm
Twinkle (-) mitochondria #2
z = 2
z = 3
z = 4
z = 5
z = 6
z = 7
500 nm

**Supplemental Figure 5. Correlative light and electron microscopy (CLEM) of axonal mitochondria with or without Twinkle in cortical pyramidal neurons *in vitro*.** Related to Figure 3. (**A**) Diagram illustrating the Correlative Light and Electron Microscopy (CLEM) analysis pipeline. Primary cortical pyramidal neurons were electroporated *ex utero* at E15.5 with iRFP (cell filler), Twinkle-Venus (mtDNA-associated protein) and a mitochondrial matrix marker (mt-DsRed) and cultured for 10DIV. Selected axons were imaged using a confocal microscope, followed by serial EM imaging. Axonal segments observed with the confocal microscope were re-identified in three-dimensionally (3D) reconstructed serial EM images. (**B**) Fluorescent images of axonal segments expressing a cell filler iRFP (blue), a mtDNA nucleoid marker Twinkle-Venus (green), and a mitochondrial matrix marker mito-DsRed (magenta). The areas marked with rectangles in the upper panels are shown at a higher magnification in the lower panels. Representative mitochondria with or without Twinkle signals are indicated by magenta or yellow arrowheads, respectively. Scale bars: 20 μm (upper panels), 10 μm (lower panels). (**C-D**) Serial electron micrographs showing the Twinkle(+) mitochondrion (C) and Twinkle(−) mitochondria (D), indicated by arrowheads in (B). Mitochondria and their cristae are clearly visible and segmented in red and cyan, respectively, in the bottom panels. See Supplementary movies S2 and S3 for an overlay of light and electron microscopy images and 3D reconstructions of mitochondria and cristae. Scale bars: 500 nm.

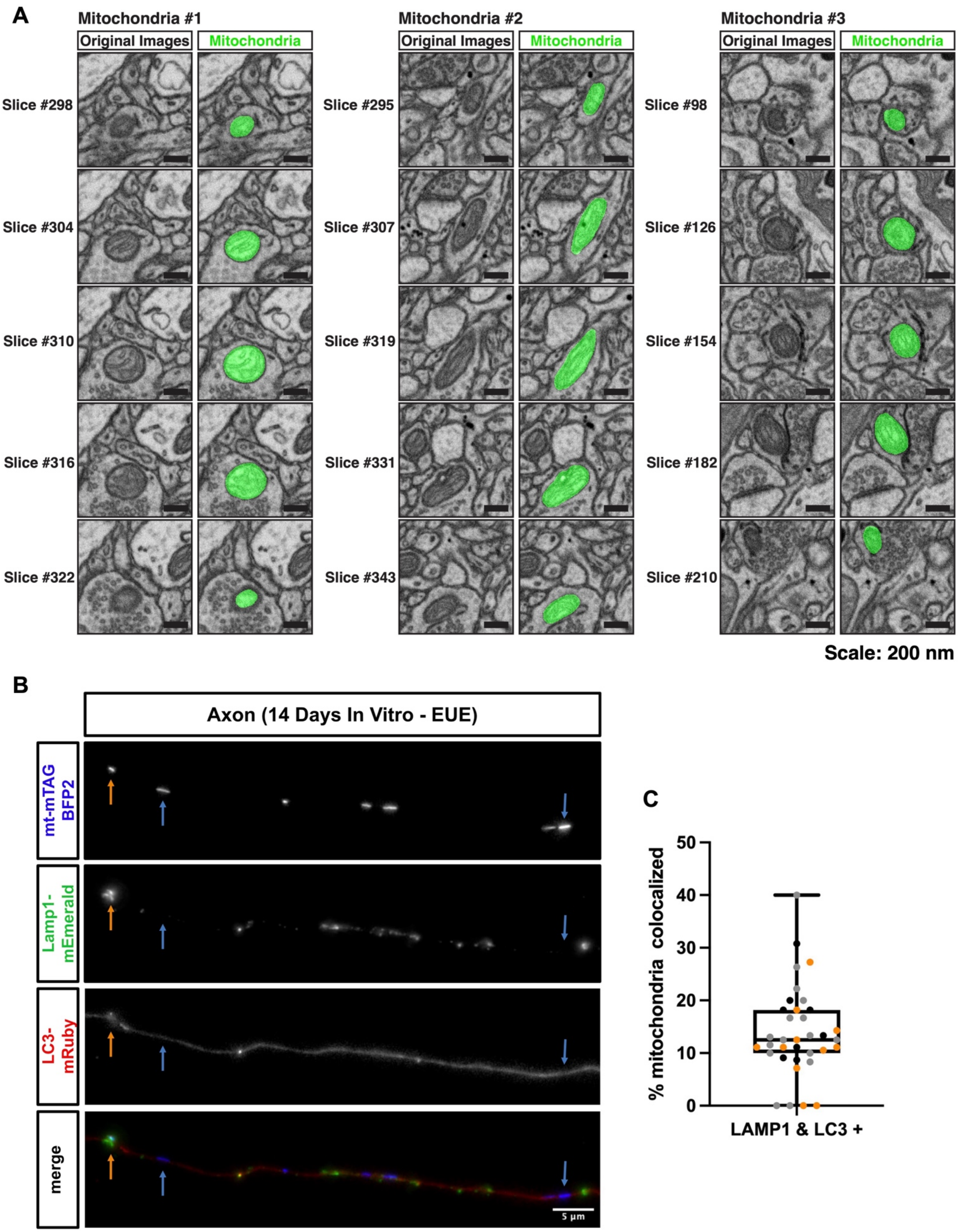
A
Mitochondria #1
Original Images
Mitochondria
Mitochondria #2
Original Images
Mitochondria
Mitochondria #3
Original Images
Mitochondria
Slice #298
Slice #304
Slice #310
Slice #316
Slice #322
Slice #295
Slice #307
Slice #319
Slice #331
Slice #343
Slice #98
Slice #126
Slice #154
Slice #182
Slice #210
Scale: 200 nm
B
Axon (14 Days In Vitro - EUE)
mt-mTAG BFP2
Lamp1-mEmerald
LC3-mRuby
merge
5 µm
C
% mitochondria colocalized
0
10
20
30
40
50
LAMP1 & LC3 +

**Supplemental Figure 6. Ultrastructural and light microscopy evidence showing that axonal mitochondria are rarely associated with autophagolysosomes.** Related to Figure 3. (**A**) Presynaptic mitochondria in adult mouse excitatory axons possess cristae *in vivo*. Three representative examples of presynaptic boutons forming synapses onto dendritic spines (excitatory glutamatergic synapses) were selected from, publicly available[39], serial electron microscopy images of mouse neocortex layer 1 (see text for detail). See Supplementary movies S4 for 3D reconstructions of the mitochondria. Scale bars: 200 nm. (**B**) Representative images of mitochondria, Lamp1 vesicles and LC3 accumulations labeled via mt-mTAGBFP2 (blue), Lamp1-mEmerald (green) and LC3-mRuby (red) in a 14 DIV cortical neuron. (**C**) Quantification of the percentage of mitochondria colocalized with Lamp1/LC3 positive structures showing that a low (~15%) percentage of mitochondria overlap. n = 36 axons, 469 mitochondria from 3 independent cultures. Scale bar: 5 microns.

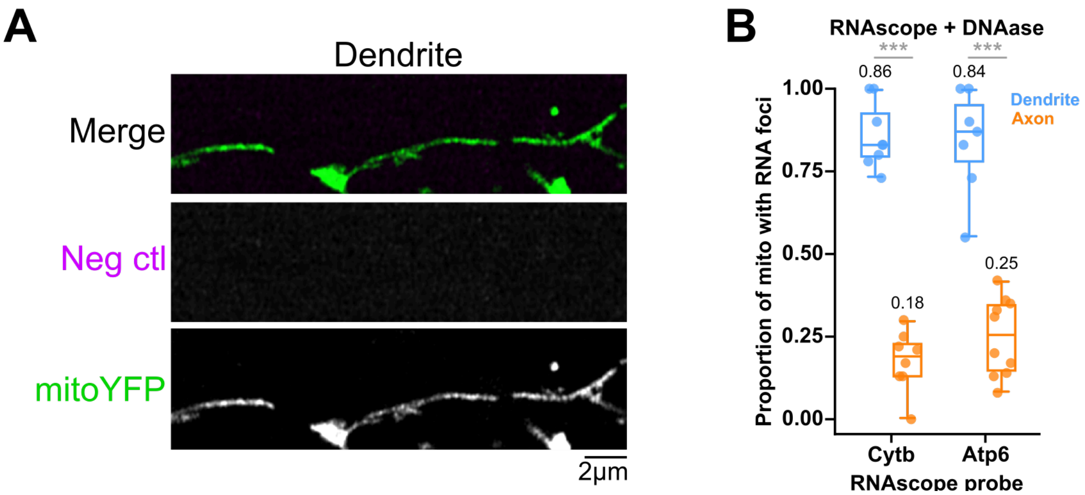


**Supplemental Figure 7. Negative controls for RNA-FISH.** Related to Figure 4. (**A**) Representative images of dendritic mitochondria (labeled with mitoYFP in green) from 21DIV neuronal cultures incubated with a non-targeting RNA-FISH probe (magenta) used as a negative control. Cultures were otherwise processed as in Figure 4h-k. (**B**) Quantification of the proportion of dendritic vs axonal mitochondria that are positive for CytB or Atp6 RNA-FISH probes. Neuronal cultures (14DIV) were treated with DNAase (100 μg/mL for 30 min at 37$^{o}$C) before incubation with RNA-FISH probes to discard unspecific binding of probes to mtDNA. Note the similarity with values shown in Fig 4l. Each dot represents individual ROIs. Number of mitochondria analyzed: Cytb dendrite n=106, Cytb axon n=113, Atp6 dendrite n=97, Atp6 axon n=107 from three independent cultures. $p<0.001$ (***) by Mann-Whitney for non-parametric data.

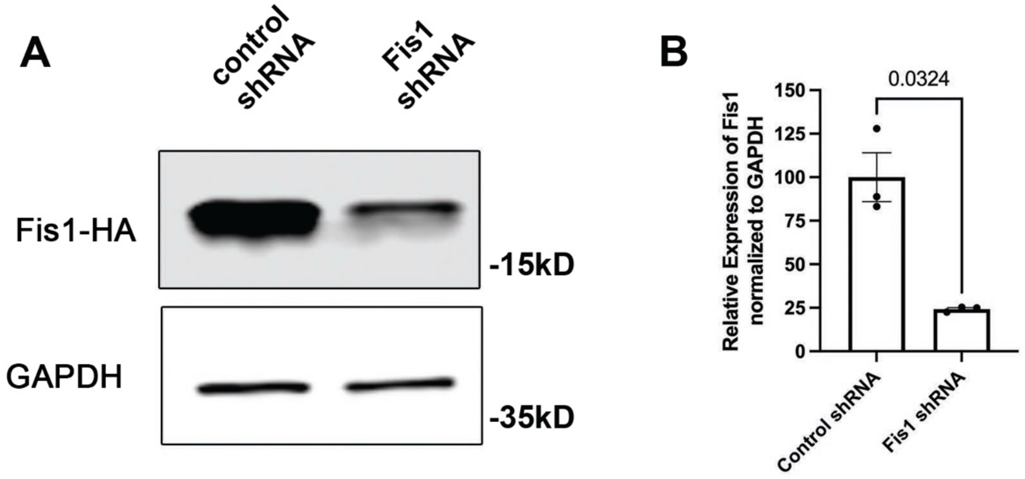


**Supplemental Figure 8. Validation of Fis1 shRNA.** Related to Figure 6.
(**A**) Representative images from western blots of HA-tagged Fis1 or loading control Gapdh following co-transfection with control shRNA (left lane) or Fis1 shRNA (right lane). (**B**) Quantification of Fis1 knockdown from 3 independent experiments. Welch's t test.

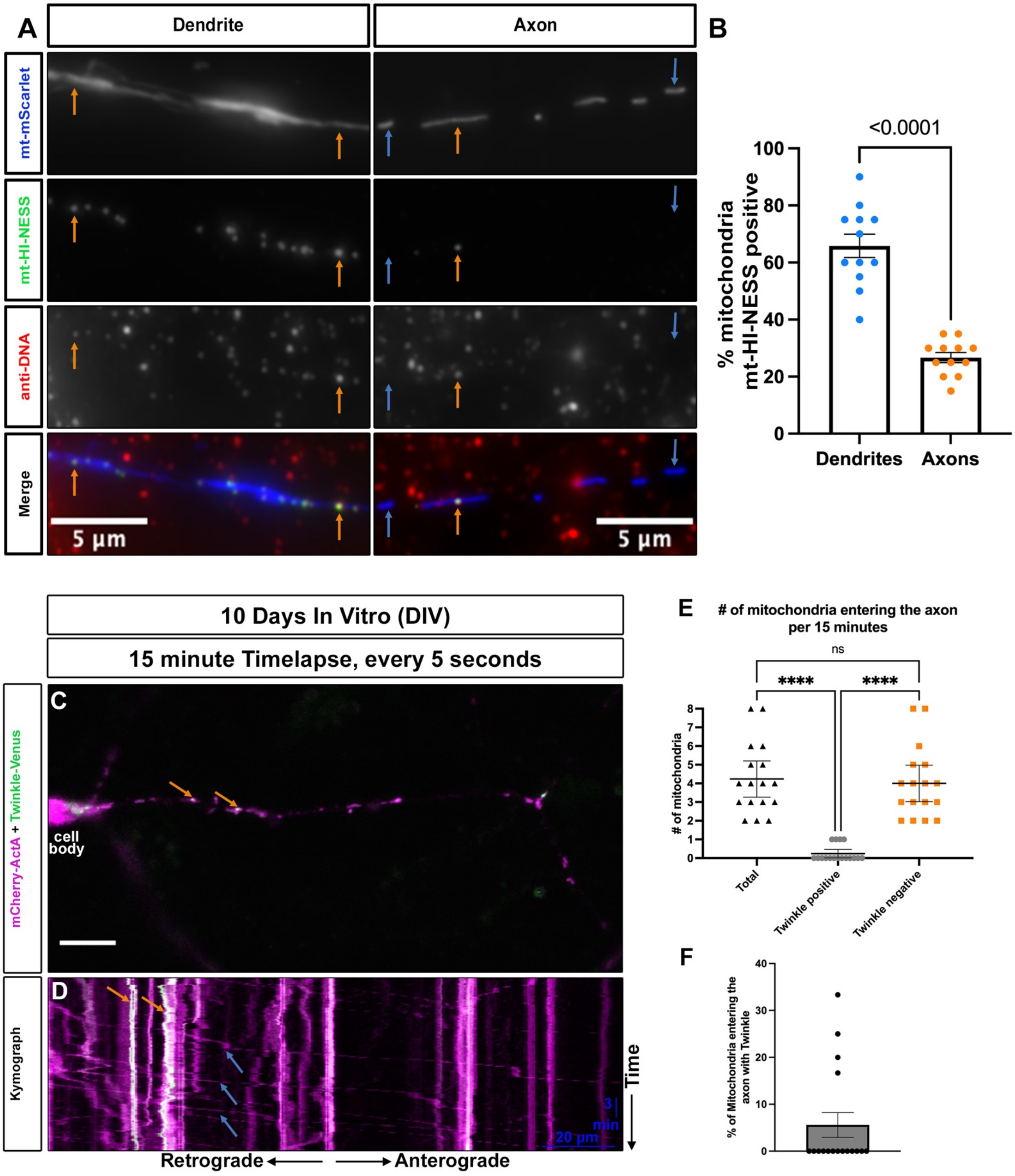

A
Dendrite
Axon
mt-mScarlet
mt-HI-NESS
anti-DNA
Merge
5 µm
5 µm
B
<0.0001
% mitochondria mt-HI-NESS positive
100
80
60
40
20
0
Dendrites
Axons
10 Days In Vitro (DIV)
15 minute Timelapse, every 5 seconds
C
mCherry-ActA + Twinkle-Venus
cell body
D
Kymograph
Time
3 min
20 µm
Retrograde
Anterograde
E
### of mitochondria entering the axon per 15 minutes
ns
****
****
### of mitochondria
Total
Twinkle positive
Twinkle negative
F
% of Mitochondria entering the axon with Twinkle
40
30
20
10
0

**Supplemental Figure 9. mt-HI-NESS reliably reports mtDNA in axonal mitochondria and monitoring of axon entry of Twinkle+ and Twinkle- mitochondria *in vitro*.** Related to Figure 6. (**A**) Representative images of dendrite (left) and axon (right) segments from 7DIV neurons that had been transfected with mt-mScarlet (blue) and mt-HI-NESS (green), followed by staining with anti-DNA (red). (**B**) Quantification of the percentage of mitochondria positive for mt-HI-NESS per segment showing that axonal mitochondria are much less likely to contain mtDNA. p value shown in the figure. Unpaired t test. n = 12 axons with 240 mitochondria, and 12 dendrites with 240 mitochondria. Blue arrows point to mt-HI-NESS negative mitochondria, while orange mitochondria point to mt-HI-NESS positive mitochondria. (**C**) Quantification of the fraction of mt-HI-NESS positive axonal mitochondria also labeled with anti-DNA immunofluorescence. (**D**) Representative image of a cell body and emerging axon of a CPN in culture (DIV10) labeled with mCherry-ActA (mitochondria) and a mtDNA-associated protein (Twinkle-Venus). (**E**) Kymograph of the axon shown in A, imaged every five seconds for fifteen minutes. (**F**) Quantification of the number of mitochondria entering the axon (left: total mitochondria), (middle: mitochondria from total with Twinkle labeling), (right: mitochondria from total without Twinkle labeling). (**G**) Quantification of the percentage of mitochondria entering the axon with Twinkle-Venus labeling, demonstrating that the majority of mitochondria entering the axon already lack markers of a nucleoid. n = 71 mitochondria from 17 axons from 3 independent cultures, ns = not significant, ****$p \leq 0.0001$ by Kruskal-Wallis test. Blue arrows point to Twinkle negative mitochondria, while orange mitochondria point to Twinkle positive mitochondria.

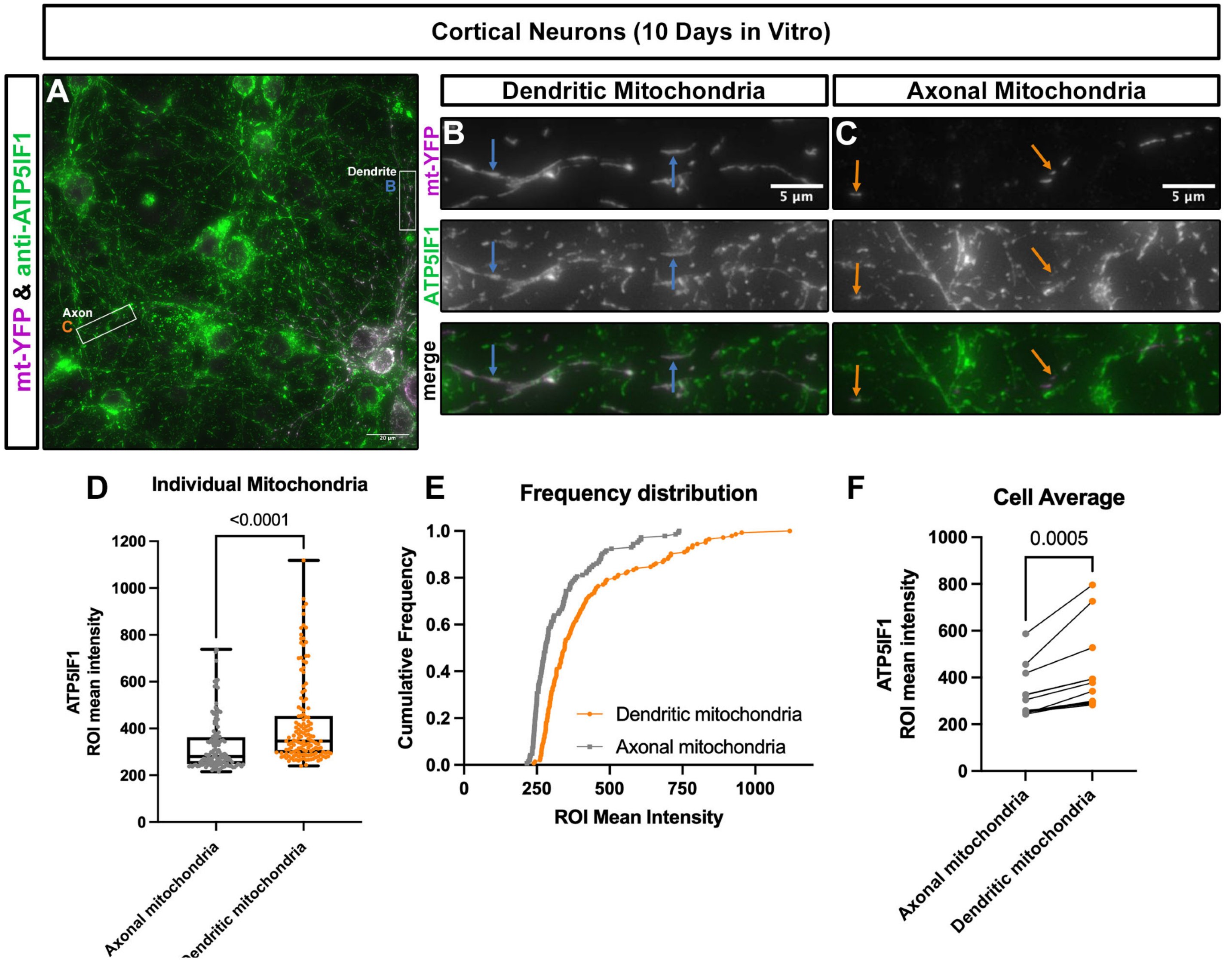


**Supplemental Figure 10. ATP5IF1 protein is expressed at significantly higher levels in dendritic than axonal mitochondria in cortical pyramidal neurons *in vitro*.** Related to Figure 7. (**A**) Representative image of 10DIV cortical neurons sparsely labeled with mt-YFP (purple) via *ex utero* electroporation and then stained with anti-Atp5if1 antibody (green). (**B**) High magnification of boxed dendritic area in A. Blue arrows point to dendritic mitochondria. (**C**) High magnification of boxes axonal region in A. Orange arrows point to axonal mitochondria. (**D**) Quantification of mean ATP5IF1 intensity in ROI drawn around mt-YFP labeled mitochondria demonstrating that ATP5IF1 protein levels are significantly higher in dendritic mitochondria than axonal mitochondria. (**E**) Cumulative frequency graph of axonal and dendritic mitochondria ATP5IF1 fluorescence intensity data showing a shift to the right for dendritic mitochondria. (**F**) Quantification of cell averages for paired axonal and dendritic ATP5IF1 mitochondrial mean intensities demonstrating higher ATP5IF1 levels in dendrites. p value in (D) via Mann-Whitney test for parametric data. p value in (F) via Wilcoxon test following a normality test. n = 144 mitochondria for both axonal and dendritic mitochondria from 12 neurons in 3 independent cultures.

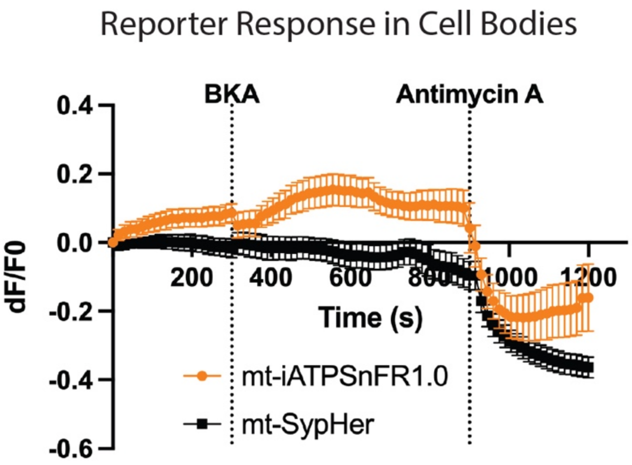


**Supplemental Figure 11. BKA treatment does not significantly alter mitochondrial pH in soma of cortical pyramidal neurons *in vitro*.** Related to Figure 7.

The soma of cortical pyramidal neurons expressing either mt-iATPSnFR1.0 (orange) or mt-SypHer (black) were timelapse imaged for 20 minutes. At 5 minutes, bath application of bongkrekic acid (BKA, an inhibitor of the ATP transporter- adenine nucleotide translocase (ANT)) at 50µM was added and fluorescence intensity of the fluorescent reporter was monitored until 15 minutes, when Antimycin A at 1.25µM was added. As expected, in the cell body (just like in dendritic mitochondria- see Figure 4), BKA addition caused an increase in mt-iATPSnFR1.0 fluorescence over the next five minutes, followed by a dramatic decrease in mt-iATPSnFR1.0 fluorescence following Antimycin A. No observable difference in mt-SypHer fluorescence was observed for the 10 minutes following BKA addition, while a decrease was observed following Antimycin A addition. This argues that the increase observed in iATPSnFR1.0 is a result of ATP increase in the matrix and not a change in pH, while most of the effect observed following Antimycin A is a result of matrix acidification. $N_{mt\text{-}iATPSnFR1.0}$ = 10 neurons from 3 independent cultures. $N_{mt\text{-}SypHer}$ = 10 neurons from 3 independent cultures.

**Supplemental Movie 1.** Related to Figure 5.

Isolation of a single axonal mitochondrion (top right corner) using a nanopipette by SICM from the culture for 7DIV. Scale bar: 5 µm.

**Supplemental Movie 2.** Related to Figure S5.

Movies corresponding to the neurons depicted in Figure S5. The plasma membrane (white) and mitochondria (magenta) of the axon from the neuron of interest were reconstructed from serial EM images. These reconstructions were overlayed with the correspond axonal segment imaged with a confocal microscope (blue).

**Supplemental Movie 3.** Related to Figure S5.

Movies corresponding to the neurons depicted in Figure S5. 3D CLEM reconstructions of mitochondria (magenta) and cristae (cyan) from a Twinkle-positive and a Twinkle-negative mitochondrion are shown. Both mitochondria contain internal cristae structures.

**Supplemental Movie 4.** Related to Figure S6.

All mitochondria (in green) reconstructed in 3D within the selected serial EM volume located at presynaptic boutons of excitatory axons, identified by synapses made onto dendritic spines. Note that each contains internal cristae structures.
